# Belayer: Modeling discrete and continuous spatial variation in gene expression from spatially resolved transcriptomics

**DOI:** 10.1101/2022.02.05.479261

**Authors:** Cong Ma, Uthsav Chitra, Shirley Zhang, Benjamin J. Raphael

## Abstract

Spatially resolved transcriptomics (SRT) technologies measure gene expression at known locations in a tissue slice, enabling the identification of spatially varying genes or cell types. Current approaches for these tasks assume either that gene expression varies continuously across a tissue or that a slice contains a small number of regions with distinct cellular composition. We propose a model for SRT data that includes both continuous and discrete spatial variation in expression, and an algorithm, Belayer, to estimate the parameters of this model from layered tissues. Belayer models gene expression as a piecewise linear function of the relative depth of a tissue layer with possible discontinuities at layer boundaries. We use conformal maps to model relative depth and derive a dynamic programming algorithm to infer layer boundaries and gene expression functions. Belayer accurately identifies tissue layers and infers biologically meaningful spatially varying genes in SRT data from brain and skin tissue samples.

## 1 Introduction

Spatially resolved transcriptomics (SRT) technologies simultaneously measure both gene expression and spatial location of cells in a two-dimensional tissue slice [12, 72, 75]. Examples of SRT technologies include in-situ hybridization (ISH) techniques such as MERFISH [22] and seqFISH+ [33], which are based on imaging fluorescence probes, and sequencing techniques such as Slide-seq [76], Slide-seqV2 [80] and the 10X Genomics Visium spatial transcriptomics platform [1, 79], which sequence barcoded mRNA molecules whose barcodes record both the spatial locations of the molecules and unique molecular identifiers (UMI) for the mRNA molecule. SRT technologies have enabled a wide range of biological analyses, including analyses on the spatial organization of different tissues [62, 11, 59, 15, 43, 32, 8, 19, 65, 30, 81, 48, 18] and inter-cellular communication [95, 20, 28, 54].

Computational analysis of SRT data presents multiple challenges. First, most SRT technologies produce high-dimensional data; for example, sequencing-based technologies such as Slide-Seq [76] and spatial transcriptomics [79] measure the expression of 10,000 to 20,000 human genes across thousands or tens of thousands of spatial locations. Second, gene expression measurements from many current SRT technologies are highly sparse, with many genes not measured even if they are expressed. For example, current sequencing-based SRT technologies have very low UMI counts per spot; e.g., in the 10X Genomics Visium platform [1] each spot has a median of about 5,000 UMI counts, while Slide-seqV2 [80] reports median UMI counts of approximately 500. Finally, jointly modeling both gene expression and the spatial location of cells in a tissue requires an appropriate model of spatial variation in gene expression, and is thus more involved than applying existing models developed for bulk or single-cell gene expression data.

Most current computational models for analyzing spatial variation in gene expression in SRT data make one of two distinct modeling assumptions. The first modeling assumption is that gene expression is determined by discrete cell types; specifically, there are a small number of clusters with distinct cell type compositions, and gene expression at a spot depends only on the cluster label, i.e. the cell type composition present at the spot. This modeling assumption is made by methods for identifying cell type clusters [99, 28, 74]. These methods implicitly address data sparsity by sharing information across nearby spots, e.g., through the use of models such as hidden Markov random fields (HMRFs). Under this modeling assumption, large differences in gene expression between clusters are allowed, which corresponds to discrete shifts in cell type composition within a tissue slice. However, methods that make this modeling assumption also assume that gene expression is constant within each cluster, and thus do not account for continuous spatial variation of gene expression within a cluster.

The second modeling assumption is that gene expression varies continuously across a tissue slice. This assumption is usually made by methods that identify spatially varying genes [83, 82, 29, 100] or that construct low-dimensional representations of measured cells [89, 86, 23]. For example, SpatialDE [83] and SPARK [82] model gene expression with a Gaussian Process in which the covariance of expression between a pair of spots decreases as their spatial distance increases. This modeling assumption is justified by the biological observation that gene expression is affected by spatial cellular environments and inter-cellular communication [53, 10, 37]. However, most methods that make this modeling assumption do not account for large discrete changes in cell type composition across the tissue and the consequent discontinuous change in expression. One exception are factor analysis approaches, such as [89, 86, 90, 23], which model gene expression as a discrete sum of continuous factors estimated from the data. These approaches can in principle model discontinuous changes in gene expression if the factors have disjoint support. However in practice these methods do not explicitly model such discontinuities. Moreover the factors learned by these methods are sometimes difficult to interpret as they are not guaranteed to correspond to distinct cell types.

We introduce a method called Belayer for analyzing SRT data using a *global* model of tissue organization and gene expression that combines both discrete and continuous spatial variation. Specifically, we define *layered tissues*, a global model of tissue organization for tissues that consist of consecutive layers of cell types. Layered tissues are common in many organs, e.g., human skin consists of three layers [49], the cerebral cortex consists of six layers [61], and the retina has ten distinct layers of neurons [60]. In the simplest case, a layered tissue has a one-dimensional spatial structure, and we model the expression of a gene as a *piecewise continuous* function of the *depth* of the tissue layers. *Piecewise* functions allow for discontinuities in expression where there are sharp changes in cell type composition in space, such as between tissue layers, while *continuous* functions model gradients of gene expression within a tissue layer, e.g. [66, 39]. To reduce overfitting with sparse SRT data, we specifically model gene expression using *piecewise linear* functions, which are specified by a small number of parameters. The inference of piecewise linear gene expression functions in 1D is related to changepoint detection [7] and segmented regression [2, 14, 93], well-studied problems in time-series analysis whose maximum likelihood solutions can be computed using dynamic programming. We extend the classical dynamic programming algorithm for segmented regression to jointly infer piecewise linear gene expression functions for all genes simultaneously. We also demonstrate that dimensionality reduction using generalized PCA (GLM-PCA) [87] preserves the one-dimensional structure of a layered tissue, thus formalizing the ad hoc dimensionality reduction steps often made when analyzing SRT data. Next, we generalize this approach to two-dimensional layered tissues by using tools from complex analysis, namely conformal maps — complex analytic functions that locally preserve angles between curves [68]— to transform a general 2D layered tissue into a layered tissue with a one-dimensional structure. We extend our dynamic programming algorithm to more general 2D layered tissues. For tissues whose layer boundaries are lines, our DP algorithm is similar to the classical Nussinov algorithm for RNA secondary structure [69].

We implement our algorithms in a method called Belayer and we apply Belayer to simulated SRT data and to three real SRT datasets, including 10X Visium data from the human dorsolateral prefrontal cortex [62] and a mouse skin wound [34] and Slide-SeqV2 data from the mouse somatosensory cortex [80]. We demonstrate that Belayer achieves higher accuracy in clustering tissue layers compared to state-of-the-art SRT clustering methods. Moreover, we also demonstrate that the piecewise linear gene expression functions learned by Belayer enable the identification of spatially varying genes, and have higher accuracy in identifying tissue-specific marker genes compared to commonly-used methods for the identification of spatially varying genes.

## 2 Results

### 2.1 Belayer Algorithm

We introduce Belayer, an algorithm for inferring spatial patterns of gene expression in spatially resolved transcriptomics (SRT) data from *layered* tissue slices. Belayer has three defining characteristics (Figure 1). **(1)** The expression of each gene is modeled as a *piecewise linear* function of the *relative depth* within each tissue layer. **(2)** *Conformal maps*, a tool from complex analysis, are used to transform the geometry of each tissue layer to that of a vertical strip and obtain the relative depth of curved tissue layers. **(3)** A dynamic programming algorithm learns tissue layers and piecewise linear gene expression functions. We describe these characteristics in more detail below.

**Figure 1:**
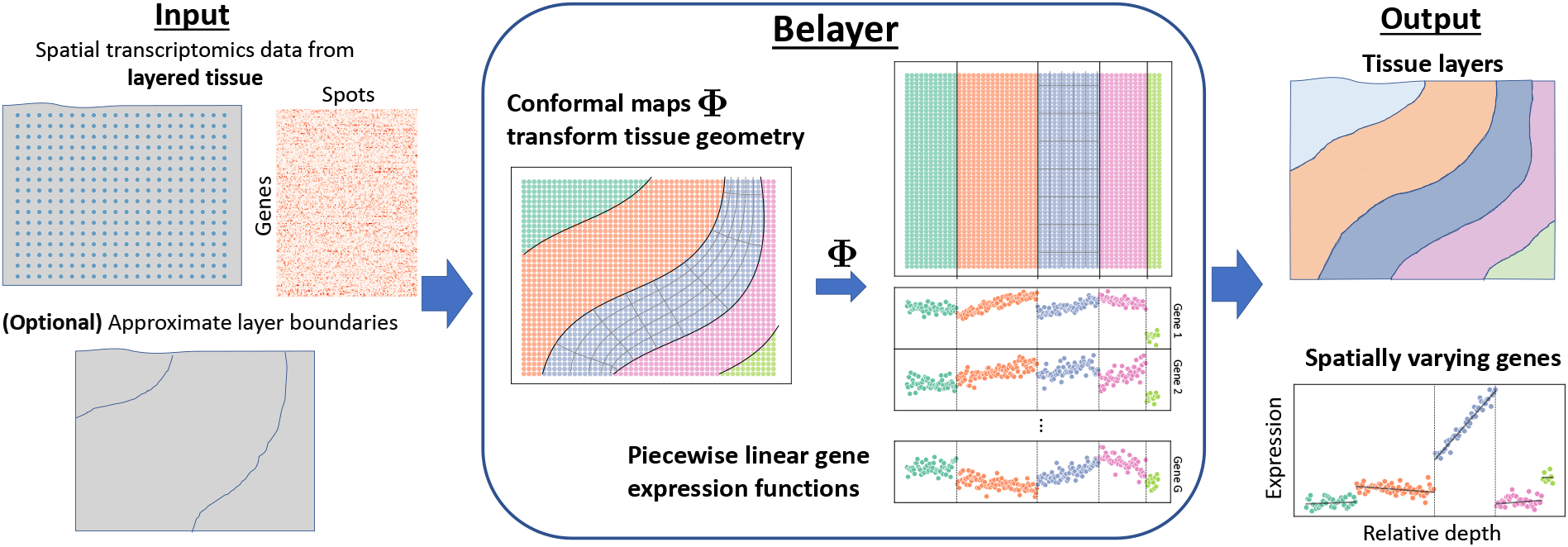
**(Left)** The input to Belayer is spatially resolved transcriptomics data from a layered tissue slice, and optionally approximate layer boundaries for the tissue slice. **(Middle)** Belayer uses conformal maps Φ = (Φ_1_,…, Φ,_L_) to transform the geometry of each tissue layer to that of a vertical strip. Belayer then models the expression of each gene in the transformed geometry as a piecewise linear function of the relative depth of the tissue layers. **(Right)** The outputs of Belayer are (1) the tissue layers, and (2) the piecewise linear gene expression functions. The latter can be used to identify spatially varying genes, such as those with large layer-specific slopes (expression gradients).

Belayer models the expression of each gene as a piecewise linear function of the spatial location. Specifically, suppose a two-dimensional tissue slice *T* consists of *L* layers *R*_1_,…, *R_L_* with a boundary curve Γ_*ℓ*_ between each pair (*R_ℓ_*, *R*_*ℓ*+1_) of adjacent layers. In the simplest case, the tissue slice *T* has “axis-aligned” layer boundaries Γ_*ℓ*_, that is layer boundaries Γ_*ℓ¿*_ that are parallel to the y-axis (Figure 7B). In this case, we model the normalized expression *f_g_*(*x, y*) of each gene *g* = 1, …, *G* at spatial location (*x, y*) ∈ *T* as a *piecewise linear* function of the x-coordinate:

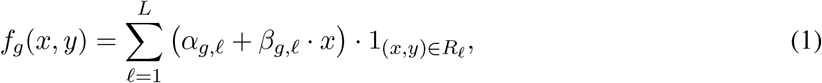

where *α_g,ℓ_* and *β_g,ℓ_* are the *y*-intercept and slope, respectively, of the expression of gene *g* in layer *R_*ℓ*_. The linear functions *a_g,ℓ_* + *β_g,ℓ_* · *x* model smooth gradients of expression within each layer *R_ℓ_*, while the pieces 1(_*x,y*_) ∈ *R_ℓ_* allow for discontinuities in expression at the layer boundaries Γ__ℓ__*. For tissues with axis-aligned layer boundaries Γ_*ℓ*_, the *x*-coordinate is the depth of position (*x, y*) in tissue layer *R_ℓ_* (Figure 7B).

**Figure 7:**
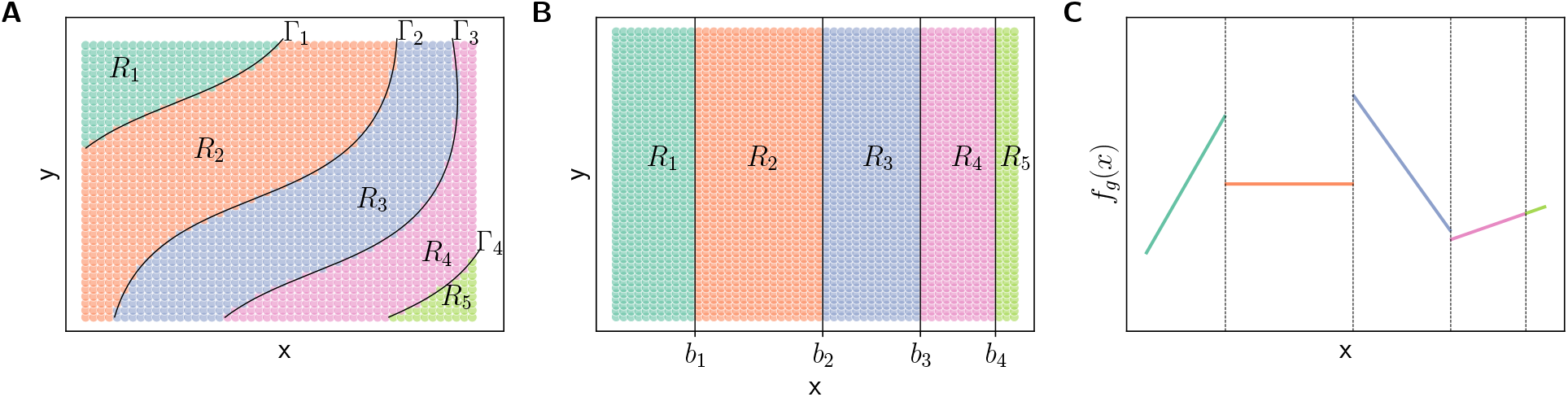
Layered tissue slices and piecewise linear expression functions. **(A)** A 5-layered tissue slice *T* with layers *R*_1_,…, *R*_5_ and layer boundaries Γ_1_,…, Γ_4_. **(B)** An axis-aligned 5-layered tissue slice with layers *R*_1_,…, *R*_5_ and layer boundaries *x* = *b*_1_,…, *x* = *b*_4_. **(C)** Expression function *f_g_* (*x*) for gene *g* along the *x* coordinate for the axis-aligned layered tissue slice in (B). For all panels, colors indicate layers and black lines indicate layer boundaries.

For a tissue slice *T* with arbitrary layer boundaries Γ_*ℓ*_, we generalize our model so that the expression *f_g_*(*x, y*) of gene *g* is a piecewise linear function of a quantity that we call the *relative* depth Φ_*ℓ*_(*x, y*) of position (*x, y*) in layer *R_ℓ_*:

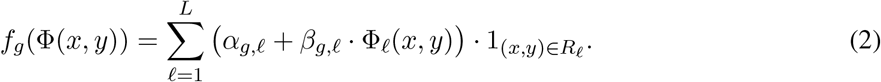

We model the relative depth Φ_*ℓ*_(*x, y*) in each layer *R_*ℓ*_* with a *conformal map*. Conformal maps are functions from the complex plane to the complex plane that locally preserve angles between curves. Conformal maps are often used in engineering and physics applications to solve differential equations with complicated boundary conditions by transforming the domain of the differential equation to a simpler geometric structure [68, 5].

Given an *N* × *G* transcript count matrix **A** = [*a_i,g_*], where *a_i,g_* is the count of gene *g* in spot *i*, and spatial coordinate matrix **S** = [**s**_*i*_] where **s**_*i*_ = (*x_i_*, *y_i_*) are the coordinates of spot *i*, Belayer aims to estimate layers 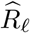, conformal maps 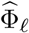, and piecewise linear functions 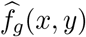 that maximize the likelihood of the observed SRT data (**A**, **S**):

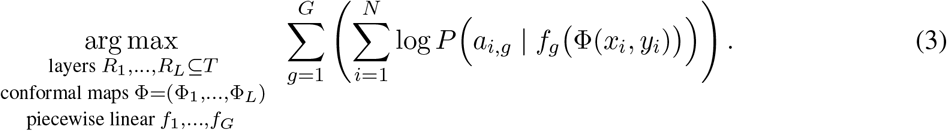

The layers 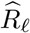 estimated by Belayer correspond to anatomical structures or other spatial partitions of the tissue slice *T* with distinct gene expression patterns. The estimated piecewise linear functions 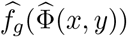 describe the spatial expression patterns of each gene g, and can be used to identify genes with potentially interesting spatial expression patterns, including genes with large (absolute) layer-specific slopes 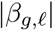 or genes *g* with large discontinuities at the boundaries 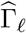 of the estimated layers 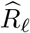. In order to easily visualize these spatial expression patterns, we combine expression values *a_i,g_* for a gene *g* from spots (*x_i_, y_i_*) with similar relative depths 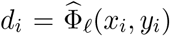 into a single “binned” expression value 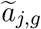 (Figure 2A). This binning procedure approximately preserves the slopes *β_g,ℓ_* and *y*-intercepts *α_g,ℓ_* of the piecewise linear gene expression functions *f_g_*(*x, y*). See Methods for more details.

**Figure 2:**
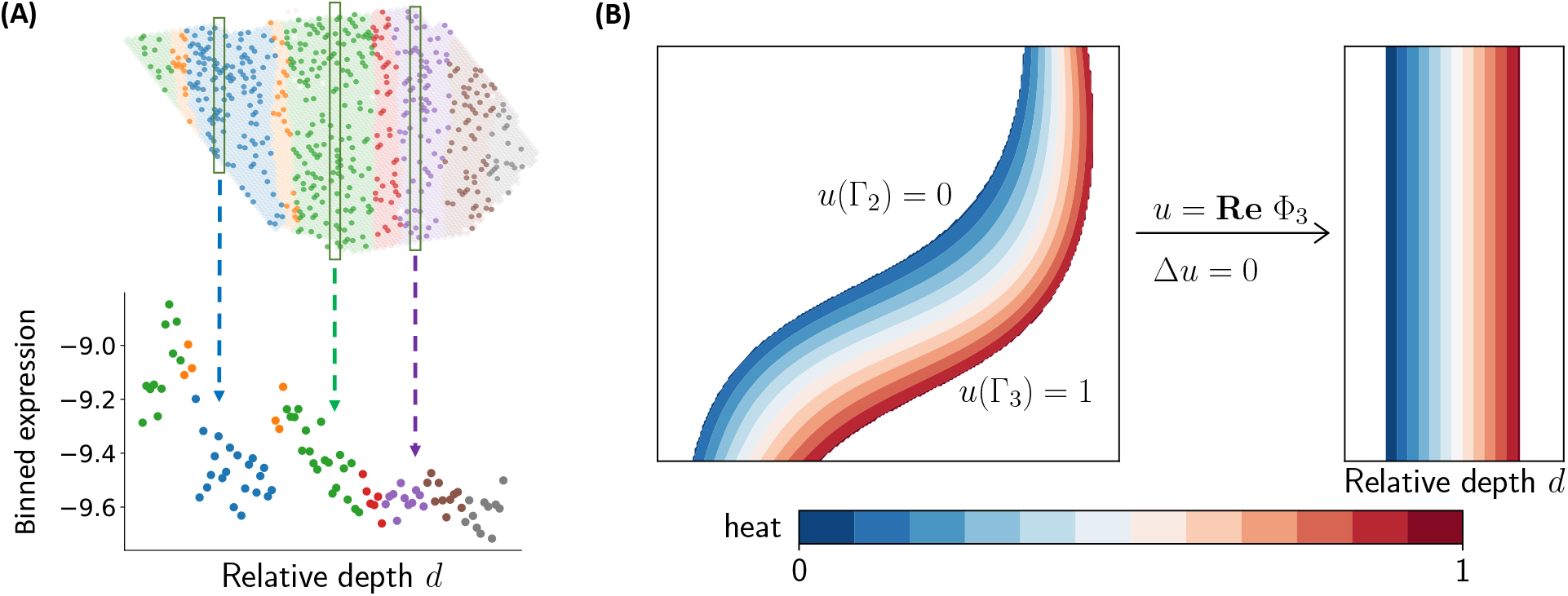
**(A)** (Top) Spatially resolved transcriptomics data from a layered tissue slice with non-zero UMIs indicated in darker color. (Bottom) Visualization of 1D expression values obtained from binning and normalization of UMI counts from 2D spots (x,y) with similar relative depths Φ_*ℓ*_(*x,y*). **(B)** The real part *u* = **Re** Φ_*ℓ*_ of a conformal map Φ_*ℓ*_ maps a curved layer to a vertical strip aligned by *x*-coordinate. *u* is a harmonic function that solves the heat equation Δ*u* = 0. The relative depth *d_i_* is the heat at spot *i* when the heat *u* = 0 is fixed at on one boundary and the heat u = 1 is fixed on the other boundary of the region. The color shows contours of the heat *u* in the layered tissue and the vertical strip.

In general, for arbitrarily shaped layer boundaries Γ_*ℓ*_, computing the maximum likelihood in (3) is challenging. We provide dynamic programming algorithms to solve two special cases of this problem which provide useful approximations on real data. First, given *approximate* layer boundaries 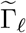 of the tissue slice *T*, such as from histological images or prior anatomical knowledge, we estimate the real part **Re**Φ_*ℓ*_ of the conformal maps Φ_*ℓ*_ by solving the heat equation (Figure 2B) with boundary conditions specified by the approximate layer boundaries 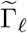. We then derive a dynamic programming algorithm for solving (3) given the estimated real parts 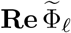 of the conformal maps Φ_*ℓ*_. Second, when the layer boundaries Γ_*ℓ*_ are *lines*, we derive another dynamic programming algorithm to compute (3) in combination with solving multiple instances of the heat equation with different boundary conditions. Our algorithm is inspired by the classical dynamic programming algorithm for segmented regression [2, 14, 93] and Nussinov’s algorithm for RNA folding [69]. We also derive a more efficient algorithm for the special case when the layer boundaries Γ_*ℓ*_ are *parallel* lines, in which case it is not necessary to solve the heat equation. See Methods for more details on these algorithms as well as our model selection procedure for choosing the number *L* of layers.

### 2.2 Evaluation on simulated data

We evaluated Belayer on two sets of simulated SRT data. In the first simulation, we generated SRT data (**A**, **S**) from a layered tissue slice *T* according to our piecewise linear model (2). In the second simulation, we used the Splatter package [96] to generate realistic single-cell SRT data from a layered tissue slice. In both simulations, we found (Figure S5A and Figure S6) that Belayer had higher accuracy in identifying the tissue layers compared to two other existing methods for analysis of SRT data, BayesSpace [99] and SpaGCN [40], and compared to analysis with SCANPY [92] which does not use spatial information. In the first simulation, where we have knowledge of the true piecewise linear gene expression functions *f_g_*(*x, y*),we also observed (Figure S5B) that Belayer accurately estimated the parameters *α_g,ℓ_, β_g,ℓ_* of each piecewise linear gene expression function *f_g_*(*x,y*). The latter evaluation demonstrates the accuracy of using the parameters estimated by Belayer in downstream tasks such as identifying spatially varying genes. See Methods for details on both simulations.

### 2.3 Dorsolateral Prefrontal Cortex

Next, we evaluated Belayer on spatially resolved transcriptomics data from the human dorsolateral prefrontal cortex (DLPFC) obtained using the 10X Visium technology [62]. This dataset consists of 12 DLPFC tissue slices from three donors. Each slice was manually annotated [62] into six layers and white matter (WM). A list of 128 marker genes for the DLPFC was obtained from [62] and previous analyses [63, 97].

#### 2.3.1 Cortical layer identification

We first analyzed the tissue layers identified by running Belayer on the DLPFC tissue slices. For the four tissue slices from Donor 1, the manually annotated layer boundaries are approximately lines. Thus, we ran Belayer with linear layer boundaries. We compared Belayer against BayesSpace [99], stLearn [74], SCANPY [92], and SpaGCN [40] on these four tissue slices. We ran each method with the same number *L* of layers as Belayer (see Methods for details on our model selection procedure). We evaluated each method by computing the ARI between the manually annotated layer labels and the layers identified by the method. Note that in some samples, the manually annotated labels correspond to two layers separated in space. We counted these as separate layers and accordingly separated any clusters of BayesSpace, stLearn, or SpaGCN that are separated in space when computing ARIs. We found (Figure 3A) that Belayer has noticeably higher ARI compared to BayesSpace, stLearn, SCANPY, and SpaGCN. Our results demonstrate that Belayer learns more biologically relevant clusters by leveraging the layered structure of the tissue. In the Appendix, we also show that Belayer outperforms these methods using other evaluation metrics (Figure S9), and that Belayer outperforms BayesSpace when BayesSpace uses its own procedure for selecting the number *L* of layers (Table S1).

**Figure 3:**
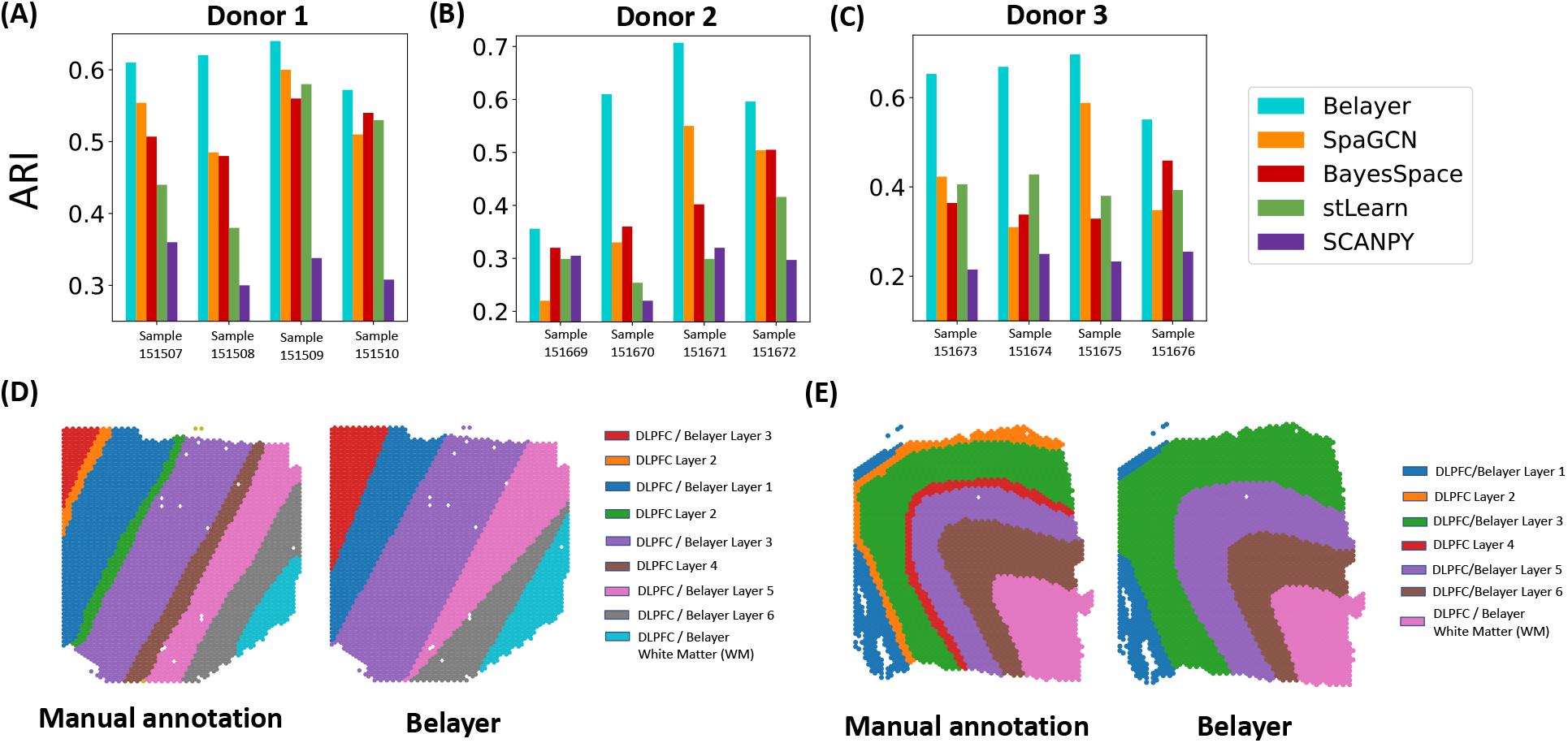
Comparison of Belayer, BayesSpace, stLearn, and SCANPY in identifying annotated layers in spatially resolved transcriptomics data from the dorsolateral prefrontal cortex (DLPFC) of three donors. **(A)** ARI for each method compared to manually annotated layers from each sample from Donor 1. **(B), (C)** Same as (A) for Donors 2 and 3. Since the tissue slices from these donors are not axis-aligned, Belayer finds conformal maps (Figure 2) to transform these tissue slices to axis-aligned tissue slices. **(D)** The layers identified by Belayer and the manually annotated layers for (axis-aligned) sample 151508 from Donor 1 and **(E)** sample 151673 from Donor 3. Layers identified by Belayer are labeled and colored according to the maximally overlapping layer from the manual annotation.

Next we compared the methods on the 8 DLPFC slices from Donors 2 and 3. The manually annotated layer boundaries of these slices are not linear. Thus, we ran Belayer with approximate layer boundaries, where we used the manually annotated layer boundaries to construct the conformal maps Φ_*ℓ*_. We found that Belayer outperformed the other methods in the identification of layers (Figure 3B-C), suggesting that our piecewise linear assumption on the expression functions is appropriate. However, we emphasize that this comparison is overly generous to Belayer since Belayer uses information from the manually annotated layer boundaries to construct the conformal maps.

#### 2.3.2 Identifying spatially varying genes

We also compared the genes with layer-specific expression patterns inferred by Belayer to the spatially varying genes identified by other methods. We derived a list of such layer-specific genes for DLPFC tissue slice sample “151508” (shown in Figure 3D) from the expression functions 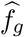 estimated by Belayer as follows. After excluding genes with low UMI counts (genes where more than 85% of spots had no UMIs), we ranked genes according to their largest layer-specific slope 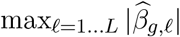 across the *L* = 6 layers 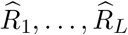 identified by Belayer. In this way, we assign high ranks to genes with large expression gradients, as gene expression gradients are known to be associated with important biological functions in the brain [42, 51, 26]. We compared the overlap in rankings between these genes and known cortical marker genes from [62, 63, 97], and performed the same comparison for ranked lists of spatially varying genes from SpatialDE [83], SPARK [82], and HotSpot [27], three methods for identifying spatially varying genes in spatially resolved transcriptomics data. We found that Belayer achieved higher AUPRC (0.032) compared to ranking genes according to the *p*-values of spatial variation computed by SpatialDE (0.017), SPARK (0.029), and HotSpot (0.029) (Figure 4A). We emphasize that all methods have low AUPRC due to the many inherent challenges of marker gene identification in SRT data. These challenges include (1) the sparsity of SRT data, (2) that the list of known marker genes are curated from multiple samples and datasets while a specific SRT sample may have variation from the “consensus”, and (3) that the list of marker genes is an incomplete representation of all genes that distinguish cortical layers.

**Figure 4:**
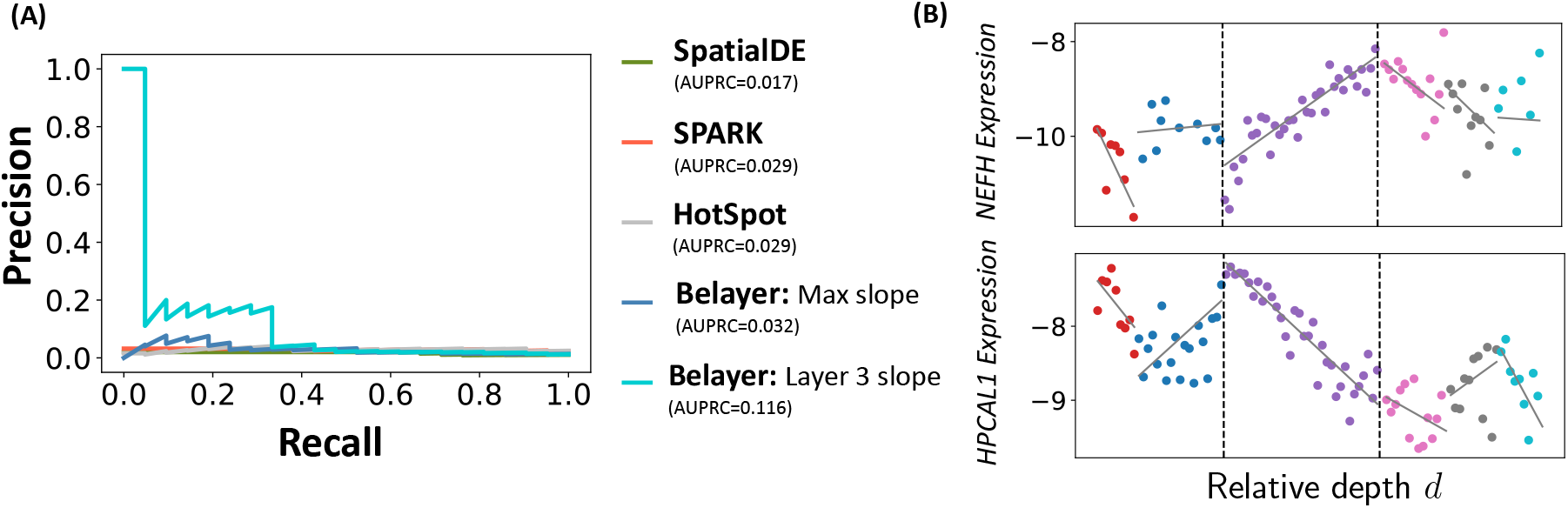
**(A)** Precision-recall curves for identifying marker genes in DLPFC sample 151508 using five different methods. “Belayer: Max slope” corresponds to ranking genes by 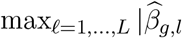 and “Belayer: Layer 3 slope” corresponds to ranking genes by 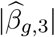. **(B)** Expression functions *f_g_*(*x*) learned by Belayer for genes *NEFH* and *HPCAL1* that have large slopes |*γ*_*g*,3_| in the third layer.

We also observed that some layers are more predictive of marker genes than others. For example, ranking genes by the magnitude of their slope 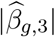 in the third layer identified by Belayer resulted in a much larger AUPRC (0.116) compared to scoring genes by their maximum slope 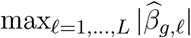 across all layers (Figure 4A). The genes *g* with large slope 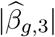 in the third layer are also biologically interesting, and we highlight two specific genes in Figure 4B. *NEFM* — the gene with the largest slope 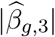 in the third layer — is a known cortical marker gene and is also a biomarker for neuronal damage [45]. On the other hand, *HPCAL1* — the gene with the fourth largest slope 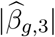 in the third layer — is not an annotated cortical marker gene but is reported to be involved in neuronal signalling [98, 91]. Our results demonstrate that incorporating layer-specific variation is important for identifying spatially varying genes, and suggest that the slopes 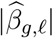 identified by Belayer are a useful criteria for identifying cortical marker genes.

We also note that the third cortical layer identified by Belayer is a combination of two manually annotated layers. Ranking genes g by their estimated slope in either one of the two manually annotated layers has smaller AUPRC compared to ranking genes by their slope in the third layer identified by Belayer. This suggests that while the layers identified by Belayer do not exactly match the manually annotated layers, they potentially correspond to other biologically relevant partitions of the tissue.

### 2.4 Mouse skin dataset during wound healing

Next, we analyzed SRT data from a mouse skin wound obtained using the 10X Visium technology [34]. Foster et al. [34] manually annotated the spots in this dataset into one of the three layers of skin: epidermis, dermis, and hypodermis. We use the sample corresponding to postoperative day 14 which contains the largest number of spots.

We evaluated Belayer’s ability to identify the 3 manually annotated layers of the skin. We used the left and right tissue boundaries as two approximate layer boundaries to estimate the conformal maps **Φ** (Figure S13A) and ran Belayer with *L* = 3 layers as determined by the model selection precedure (Figure S7). We compared the *L* = 3 layers estimated by Belayer with the *L* = 3 clusters of BayesSpace, stLearn, SCANPY, and SpaGCN. We observe (Figure 5A) that stLearn and Belayer achieve a higher ARI compared to other methods. Moreover, while stLearn obtains a slightly higher ARI than Belayer ARI = 0.564 for stLearn, ARI = 0.520 for Belayer), stLearn predicts a discontinuous epidermal layer, which is inconsistent with the manual annotation (Figure 5B). By leveraging the layered geometry of the tissue, Belayer accurately identifies biologically meaningful skin layers.

**Figure 5:**
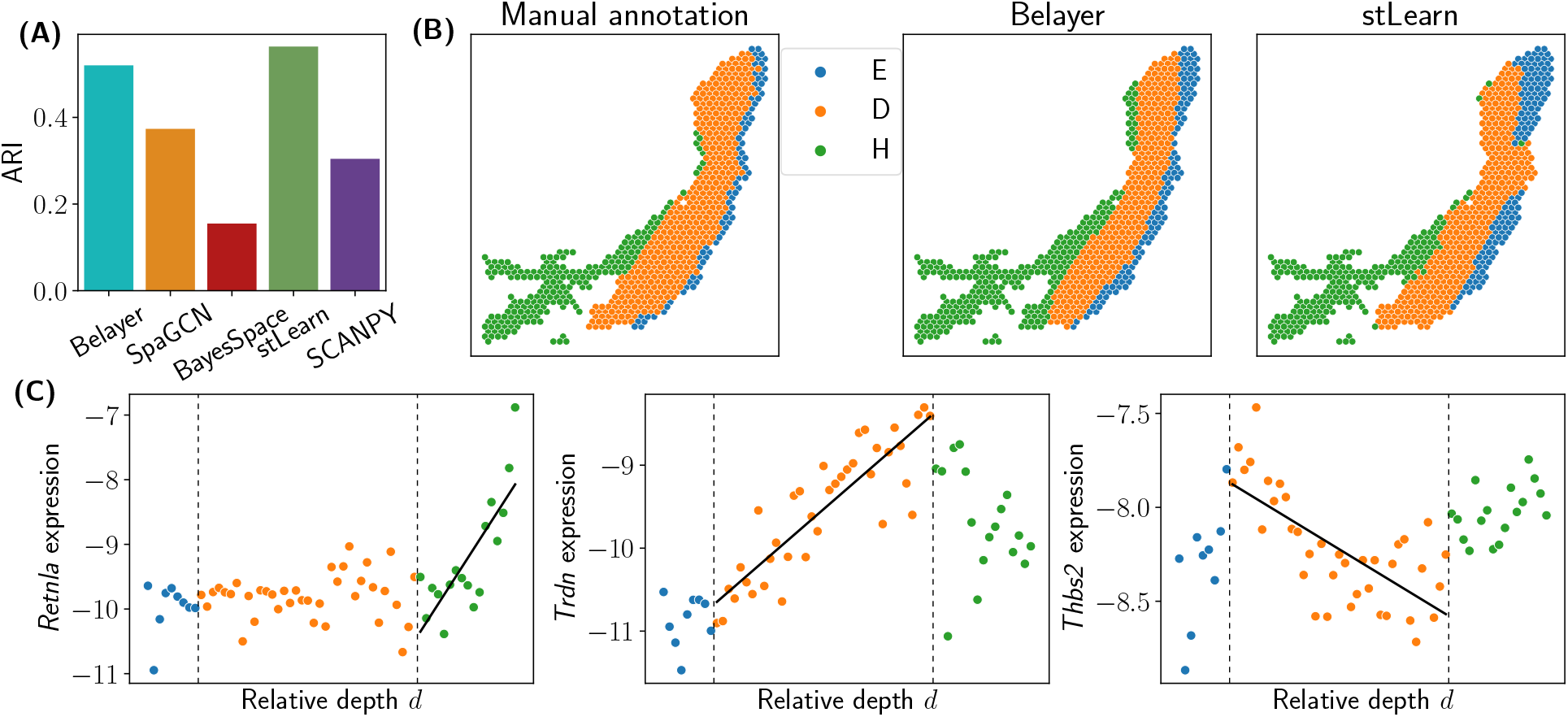
(A) Comparison of Belayer, BayesSpace, stLearn, SCANPY, and SpaGCN in identifying skin layers in a mouse skin Visium sample. (B) Manually annotated layers and layers identified by Belayer and stLearn. In the legend of manual annotation, “E” indicates epidermis, “D” indicates dermis, and “H” indicates hypodermis. (C) Expression functions learned by Belayer for three genes with large slopes 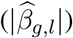 Dashed vertical lines indicate layer boundaries identified by Belayer. Solid lines show the expression function for the layer where the gene has a large slope.

We also analyze the genes *g* that Belayer infers to have large positive or negative slopes |Γ_*g,ℓ*_| (Figure 5B). We find that Belayer recapitulates some skin wound marker genes reported in [34]. These include: *Thbs2*, a gene involved in mouse cutaneous wound healing [3] that has the 8th most negative slope 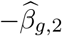 in the dermis layer (Figure 5C), and *Fn1*, a marker gene for a fibroblast cluster [34] that has the 11th largest slope 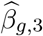 in the hypodermis layer.

Belayer also identifies genes related to the wound healing process which were not *reported in* [34] (Figure 5C). For example, *Retnla* has the largest positive slope 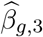 in the hypodermis layer, and it encodes a protein that is known to be an effector molecule in the skin wound healing process [46]. *Trdn* has the largest positive slope 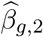 in the dermis layer, and it encodes an integral transmembrane protein involved in muscle contraction [70], a healing response for skin wounds.

### 2.5 Somatosensory Cortex

We evaluated Belayer on SRT data for a single slice of the somatosensory cortex [19] obtained using the Slide-SeqV2 technology [80]. The somatosensory cortex is a part of the neocortex and consists of six layers [44]. We obtained a list of 30 marker genes for the somatosensory cortex from [25, 64] that are also measured in this dataset.

We ran Belayer with two approximate layer boundaries, the top and bottom boundaries of the tissue slice which were chosen from visual inspection of the RCTD layer annotations [19] (Figure S12). We compared the *L* = 5 layers identified by Belayer to the clusters identified by SpaGCN [40], SCANPY, and stLearn. We evaluated the accuracy of each method according to the cell types annotated by RCTD [19], which integrated the Slide-SeqV2 data with a reference scRNA-seq dataset. We do not compare to BayesSpace [99], since the current implementation of this method requires SRT data obtained from the ST or 10X Visium platforms. We find that compared to SpaGCN, Belayer distinctly identifies the layers of the somatosensory cortex (Figure 6A). We also observe that the layers identified Belayer are more similar to the RCTD cell type annotations than the clusters identified by SpaGCN, stLearn, and SCANPY (Figure S11A). Moreover, compared to the RCTD cell type annotations — which are obtained through a reference scRNA-seq dataset with cell type annotations — we find that the layers identified by Belayer correspond more closely to the distinct layers of the somatosensory cortex. In particular, Belayer is able to clearly identify the L2/3 layer (Figure 6A) while RCTD models this layer as a mixture of different cell types from layers 2 through 6, which is biologically inconsistent with the layered geometry of the somatosensory cortex [44]. These results demonstrate that Belayer more accurately learns tissue layers by utilizing a global model of layered tissues.

**Figure 6:**
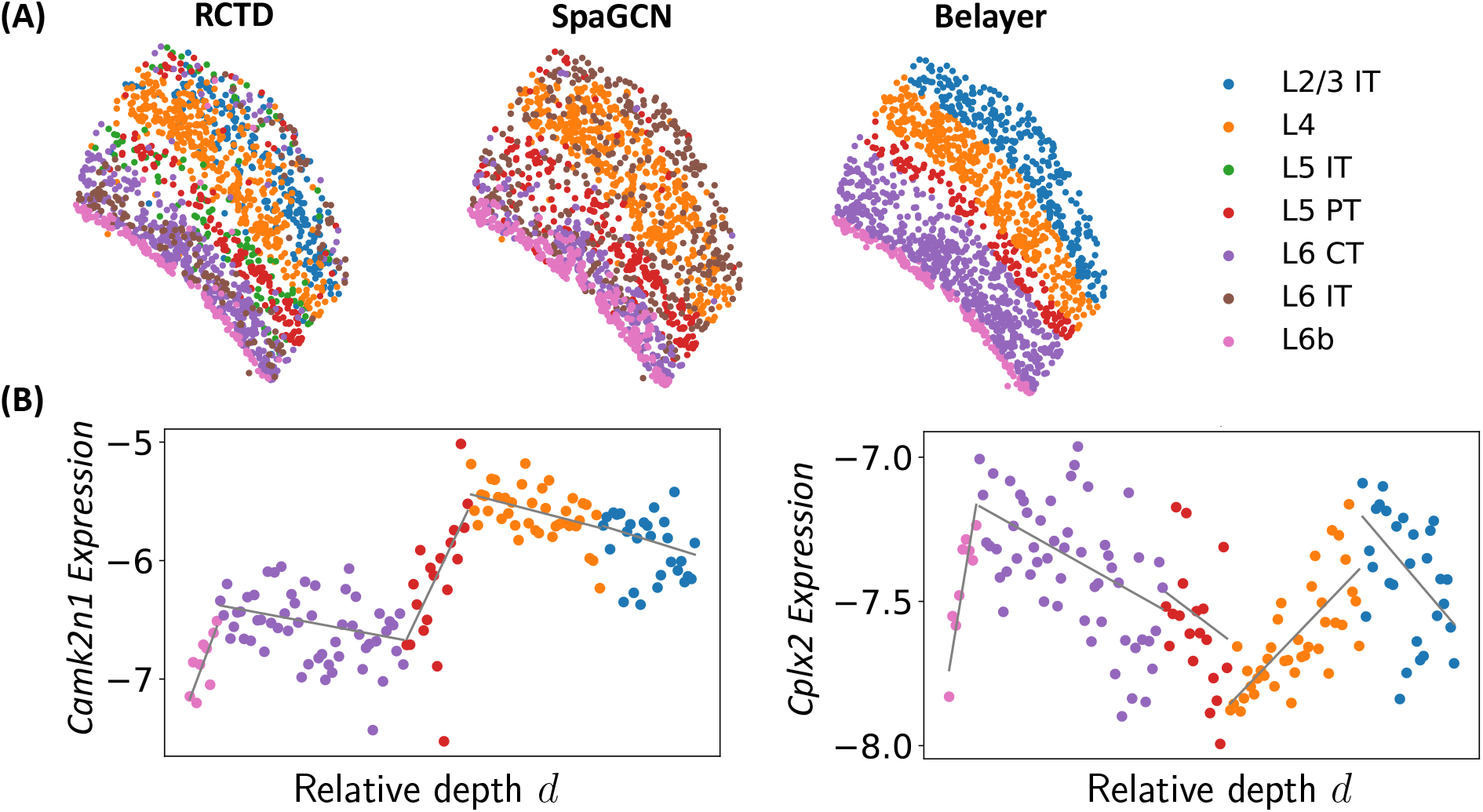
(A) Layers identified by RCTD, SpaGCN, and Belayer in a somatosensory cortex Slide-SeqV2 sample. The layers identified by SpaGCN and Belayer are labeled and colored according to the maximally overlapping layer from the RCTD cell type annotations. (B) Expression function learned by Belayer for genes *g* with large slopes 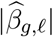.

We also compared the genes with layer-specific expression patterns inferred by Belayer to the spatially varying genes identified by SpatialDE [83], SPARK [82], and HotSpot [27] following the same procedure as in Section 2.3.2. We found (Figure S11B) that Belayer achieved higher AUPRC (0.06) compared to ranking genes by the *p*-values of spatial variation computed by SpatialDE (0.024) and SPARK (0.049), and had comparable AUPRC compared to HotSpot (0.061). Moreover, if we rank genes according to the magnitude of their slope 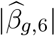 in the layer 2/3 intratelencephalic region estimated by Belayer (“L2/3 IT” in Figure 6A), then Belayer has larger AUPRC (0.099) compared to SpatialDE, SPARK, and HotSpot. As in Section 2.3.2, we emphasize that all methods have low AUPRC because of the challenges in identifying marker genes in SRT data. In particular, the list of marker genes for the somatosensory cortex that we use for evaluation is most likely incomplete as it contains less than 30 genes, while more than 1,000 genes are estimated to be involved in neuronal functions in the cerebral cortex [97].

We highlight two genes with large layer-specific slope 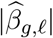 that are not in the list of marker genes but are biologically interesting (Figure 6B). *Camk2n1* has the fourth-largest layer-specific slope, and is reported to have spatiotemporal patterns of regulation [56]. *Cplx2* has the tenth-largest layer-specific slope, and mutations in this gene are reported to contribute to cognitive dysfunction in schizophrenia [38]. Our results demonstrate that the spatial patterns identified by Belayer are important in identifying biologically relevant genes.

## 3 Discussion

We introduce a new method, Belayer, to analyze spatial variation in gene expression from spatially resolved transcriptomics (SRT) data from *layered tissues*. Belayer models the expression of each gene with a *piecewise linear function* of the relative depth of the tissue layers. This piecewise linear model allows for both discrete changes in expression between layers – e.g., due to changes in cell type compositions – as well as continuous variation in gene expression within layers – e.g., due to gradients of gene expression. In the simplest case of an axis-aligned tissue structure, we infer the maximum likelihood expression function using a dynamic programming algorithm that is related to the classical problems of changepoint detection [7] and segmented regression [2, 14, 93]. We extend our approach to arbitrary layered tissues using the theory of conformal maps [68] from complex analysis [4], which are related to harmonic functions – and more specifically the heat equation – and are often used to solve partial differential equations in engineering applications. We provide algorithms to solve two important special cases of the general problem for linear layer boundaries and pre-specified approximate layer boundaries.

We show that Belayer outperforms existing approaches in identifying the tissue layers and spatially varying genes in both simulated SRT data and three real SRT datasets: 10X Visium data from the human dorsolateral prefrontal cortex (DLPFC) [62], Slide-SeqV2 data from the mouse somatosensory cortex [80], and 10X Visium data from a mouse skin wound [34]. Additionally, Belayer discovers genes with layer-specific and continuous spatial variation in expression that correspond both to known tissue-specific marker genes and genes with potentially novel tissue-specific functions. These results demonstrate that our piecewise linear model is a reasonable approach for the identification of tissue layers and the discovery of layer-specific marker genes.

There are a number of directions for future investigation. The first direction is to further investigate the expression functions learned by Belayer. For example, the observation that the slopes of the expression functions learned by Belayer recovers known tissue-specific marker genes better than existing methods inspires further study of the novel genes that are identified by our model, including genes with discontinuities and changes in slope at layer boundaries. It would also be of interest to relate the slopes of the gene expression functions to biological quantities such as the cell type proportion at different spots, which could be estimated using SRT deconvolution methods [19, 78, 9, 47, 31]. Second, it would be desirable to provide an algorithm to solve (or approximately solve) the maximum likelihood problem in (3) for arbitrary layer boundaries. One possible approach is to extend the dynamic programming algorithm for lines to a larger class of layer boundaries. Third, it would helpful to further minimize overfitting by incorporating regularization and rigorous statistical testing into our algorithm; e.g., using the Chow test [24] for change-point detection. Such extensions might also allow for nonlinear piecewise continuous expression functions, assuming the data has sufficient spatial resolution. The fourth direction is to extend the definition of a layered tissue to account for more complex tissue geometries such as muscle tissues with concentric or striated layer structures or 3-D tissues [6]. While the heat equation can be solved over closed domains or in three dimensions, deriving an appropriate representation of piecewise constant/linear expression functions for more complicated geometries requires further work. In addition, the problem of inferring the layers and the conformal map from more complicated geometries may not be straightforward. We also note that while some tissues may not have a layered structure, they may be subdivided into layered regions. For example, the mouse neocortex contains a layered somatosensory cortex, which are segmented using prior biological knowledge [19]. It would be useful to extend our definition of layered tissues towards these “piecewise” layered tissues as well as to systematically identify these layered tissue regions, perhaps by first coarsely subdividing a tissue using other SRT analysis methods. Finally, it would be interesting to extend our approach to model other spatially resolved data, including SRT data obtained from ISH technologies [57, 73] and spatial proteomics [58].

## Methods

### 4 Layered tissues and expression functions

Modeling spatial patterns of gene expression is complicated by the large variability in the spatial structure of tissues — e.g. some parts of brain tissues have a layered structure while muscle tissues have a striated structure [6]— as well as the sparsity of the gene expression profiles produced by current spatially resolved transcriptomics (SRT) technologies. To avoid overfitting the data, it is helpful to make simplifying assumptions about the spatial structure of the tissue slice T and/or the spatial patterns of gene expression. Here, motivated by the layered structure of the skin, brain, eyes, and other organs [6], we focus on *layered tissue slices* (Figure 7A) which we define as follows.

#### Definition 1.

*An L*-layered tissue slice *is a region* 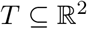 *containing L* – 1 *non-intersecting smooth curves* Γ_1_,…, *Γ_*L*–1_, *or layer boundaries, satisfying**:

1. *each curve* Γ_*ℓ*_ *has end points on the boundary ∂T* of *T*;
2. *every point p* ∈ *T is contained in a region R bounded by ∂T and* at most two *curves* Γ_*ℓ*_ *and* Γ_*ℓ*′_.

The *L* – 1 layer boundaries Γ_1_,… Γ_*L*–1_ partition the tissue slice *T* into *L* regions *R*_1_, …, *R_L_*, or *layers*. The layers *R*_1_,…, *R_L_* of a *L*-layered tissue slice *T* represent biologically distinct regions in the tissue slice *T*. For example, in a tissue slice *T* from the skin, *R*_1_, *R*_2_, *R*_3_ may represent the epidermis, dermis, and hypodermis layers, where each layer consists of unique cell types and has unique functions [49]. Without loss of generality we assume that the layers *R*_1_,…, *R_L_* and layer boundaries Γ_1_,…, Γ_*L*–1_ are ordered so that layer *R*_1_ is bounded by tissue boundary *∂T* and layer boundary Γ_1_, layer *R_*ℓ*_* is bounded by layer boundaries Γ_*ℓ*–1_ and Γ_*ℓ*_ for *ℓ* = 2,…, *L* – 1, and layer *R_L_* is bounded by layer boundary Γ_*L*–1_ and tissue boundary *∂T*.

We describe the spatial distribution of the expression of a gene *g* in a tissue *T* with an *expression function* 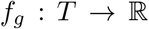, where *f_g_*(*x,y*) measures the expression of gene *g* at spatial location (*x,y*) ∈*T*. For example, a gene *g* whose expression is uniform across the tissue has a constant expression function *f_g_*(*x,y*) = *c*, while a marker gene *g* for a specific region *R* ⊆ *T* could have the expression function *f_g_*(*x, y*) = *c* · 1_{(x,y)∈*R*}_ + *c*′ ·1_{(*x,y*)∉*R*_}.

More generally, if *T* is a *L*-layered tissue slice with layers *R*_1_,…, *R_L_* and the expression of gene *g* in layer *R_ℓ_* is given by layer-specific expression function *f_g,ℓ_*(*x, y*), then gene *g* has the expression function 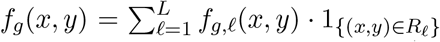. We assume that the layer-specific expression functions *f_g,ℓ_*(*x, y*) are continuous functions, and thus *f_g_*(*x, y*) is a piecewise-continuous function with discontinuities allowed at the layer boundaries Γ_*ℓ*_. These discontinuities correspond to differences in expression due to changes in cell type composition between different layers *R_ℓ_* of the tissue *T*. In the next section, we describe how one models SRT data using the expression functions *f_g_*(*x, y*).

### 5 Axis-aligned layered tissues

We begin by studying the simplest *L*-layered tissue slice: the *axis-aligned L*-layered tissue slice, where each layer boundary Γ_*ℓ*_ is a line *x* = *b_ℓ_* parallel to the *y*-axis (Figure 7B). We assume that the expression of a gene *g* at position (*x, y*) depends only on the *layer depth*, or the distance from (*x, y*) to the nearest layer boundaries *x* = *b_ℓ_*. Under this assumption, the layer-specific expression functions *f_g,ℓ_*(*x, y*) are functions of only the *x*-coordinate: *f_g,ℓ_*(*x, y*) = *f_g,ℓ_*(*x*). Thus, the expression function *f_g_*(*x, y*) also only depends on the *x*-coordinate, i.e. 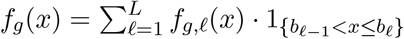 where for convenience we define *b*_0_ = –∞and *b_L_* = ∞. We call each *b_ℓ_* a *breakpoint* of the expression function *f_g_*.

The simplest model for *f_g_*(*x*) is a piecewise constant function, which corresponds to each gene *g* having a constant expression in each layer *R_ℓ_*. Such a piecewise constant model is implicit in methods that assume constant expression in contiguous regions of cell types; e.g. methods that use hidden Markov random field (HMRF) models [99, 94, 28, 85]. We generalize these approaches by modeling continuous variation in expression within a layer; e.g. due to gradients of gene expression [66, 39, 17, 21]. Since current SRT technologies have limited dynamic range and spatial resolution, inference of complicated expression functions may be prone to overfitting. Thus, we use the simplest form of continuous, non-constant spatial variation within each layer and model each layer-wise expression function *f_g,ℓ_*(*x*) as a *linear function*. Then each expression function *f_g_*(*x*) is a piecewise linear function (Figure 7C) with shared breakpoints *b*_1_,…, *b*_*L*–1_ across all genes corresponding to the axis-aligned layer boundaries Γ_1_,…, Γ_*L*–1_, i.e.

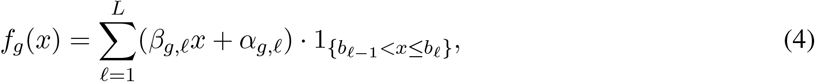

where *β_g,ℓ_* and *α_g,ℓ_* are the slope and *y*-intercept, respectively, of gene *g* in layer *R_ℓ_*. We define 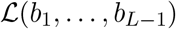 to be the set of piecewise linear functions *f*(*x*) with breakpoints *b*_1_,…, *b*_*L*ℓ1_, and we define 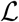 to be the set of linear functions *f*(*x*).

We contrast our model for the expression functions *f_g_* with another commonly used model for expression functions, Gaussian Processes (GPs) [83, 82, 86, 55]. In this approach each expression function 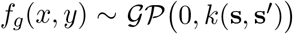 is an independent sample from a GP with mean function 0 and covariance function k(**s**, **s**′) between spatial locations **s** = (*x, y*) and **s**′ = (*x*′, *y*′). The covariance functions k(**s**, **s**′) used by existing methods, including SpatialDE [83] and SPARK [82], generate continuous functions and thus do not model piecewise continuous expression functions *f_g_* (x). We show in the Appendix that it is possible to model piecewise linear functions with GPs using a one-dimensional blockwise covariance function; however, to our knowledge such covariance functions have not been used to model SRT data.

#### 5.1 Axis-Aligned *L*-Layered Problem

We aim to infer the layer boundaries *x* = *b_ℓ_* and expression functions *f_g_*(*x*) that maximize the likelihood of the observed SRT data. We represent SRT data as an expression matrix 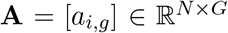 and a spatial location matrix 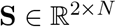 with column **s**_*i*_ = (*x_i_*, *y_i_*) indicating the spatial location of spot *i*. We define the Axis-Aligned *L*-Layered Problem as the the following maximum likelihood estimation problem.

##### Axis-Aligned *L*-Layered Problem

*Given SRT data* (**A, S**) *and a number L of layers, find layer boundaries x* = *b*_1_,…, *x* = *b*_*L*–1_ *and piecewise linear expression functions* 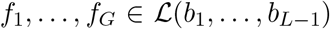 *that maximize the log-likelihood of the data*:

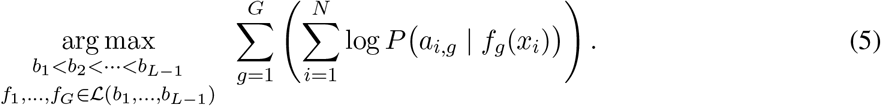

When there is *L* = 1 layer, each expression function *f_g_*(*x*) is a linear function, and thus the maximum log-likelihood (5) in the Axis-Aligned *L*-Layered Problem reduces to

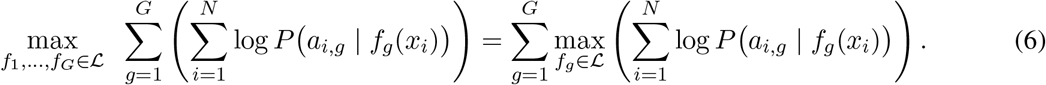

The maximization on the right-hand side of (6) is a regression problem of finding the linear function *f_g_* that best fits the observed data. Thus, (6) is solved by computing a separate regression for each gene *g* = 1,…, *G*.

More generally, when each expression function 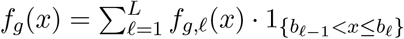 is an *L*-piecewise linear function with *known* breakpoints *b*_1_,…, *b*_*L*–1_, then the maximum log-likelihood (5) in the Axis-Aligned *L*-Layered Problem reduces to

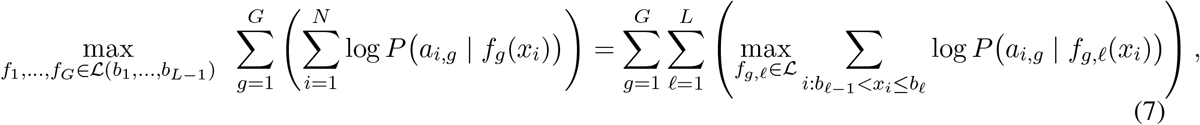

Then, each expression function *f_g_* is computed by solving *L* separate regression problems, one regression problem for each pair *x* = *b*_*ℓ*–1_, *x* = *b_*ℓ*_ of consecutive layer boundaries. On the other hand, when the breakpoints *b*_1_,…, *b*_*L*–1_* of the piecewise linear expression functions *f_g_* are *unknown*, one will have to compute a regression over *all* pairs of possible layer boundaries for each gene *g*, as we describe in the next section.

We emphasize that the regression approach described above can be used with different probability distributions *P*{*a_i,g_* | *f_g_*(*x_i_*) for the observed counts *a_i,g_*. In particular, following [87, 77, 71] we model UMI counts *a_i,g_* with a Poisson distribution 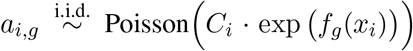 where *C_i_* is the total UMI count at spot *i*. Using the Poisson model we solve the regression problems in (7) with Poisson regression. Another alternative is to model normalized expression values *a_i,g_*, with a Gaussian distribution 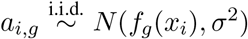 where *σ*^2^ is a shared variance parameter. Under the Gaussian model, the regression problems in (7) are linear regressions.

##### Visualization and binning of SRT data

To simplify the visualization of SRT data (**A**, **S**) from an axis-aligned tissue slice, we combine expression values for a gene from spots with the same *x*-coordinate into a single “binned” expression value (Figure 2A). Specifically, for each gene *g* we construct a binned expression value 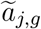 that estimates the gene expression for all spots with the same *x*-coordinate 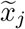. For the Poisson model the binned expression 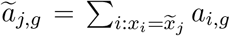 is the sum of the UMI counts *a_i,g_* at spots ***s***_*i*_ = (*x_i_, y_i_*) with *x*-coordinate 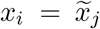, while for the Gaussian model the binned expression 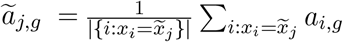 is the average expression. We show in the Appendix that binning the data does not affect inference of the expression functions *f*_1_,…, *f_G_* in the Axis-Aligned *L*-Layered Problem, as the expression functions obtained by maximizing the log-likelihood (5) with the binned expression 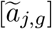 are equal to the expression functions obtained by maximizing (5) with the original expression [*a_i,g_*].

In the applications presented below, it is often the case that there are spots **s**_*i*_ = (*x_i_, *y_i_*) with *approximately* equal *x*-coordinates *x_i_*. Thus, we extend the binning approach to construct binned expression 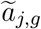 from spots **s**_i_* = (*x_i_*, *y_i_*) with *x*-coordinates 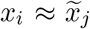. Further, for UMI count data we plot the *normalized* binned expression 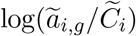, where 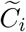 is the sum of the total UMI counts *C_i_* for all spots *i* in bin *j* (Figure 2A). These normalized counts have the same scale as the expression functions *f_g_*. See the Appendix for more details.

### 5.2 A dynamic programming algorithm for the Axis-Aligned *L*-Layered Problem

When there is only one gene, i.e. *G* = 1, then the Axis-Aligned *L*-Layered Problem is also known as *segmented regression*. Segmented regression is a classical problem in statistics and time-series analysis and has a well-known dynamic programming (DP) solution [13, 2, 14, 93]. Here, we present a DP algorithm for the Axis-Aligned L-Layered Problem that extends the classical DP algorithm for segmented regression [2, 14, 93] to any number *G* of genes.

Briefly, the DP algorithm is as follows. Let *M_n,ℓ_* be the maximum log-likelihood of the first n datapoints fit to ŕ-piecewise linear expression functions *f*_1_,…, *f_G_*, i.e.

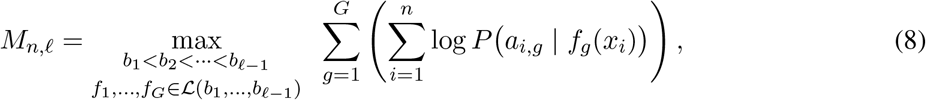

where we assume without loss of generality that the spots are ordered by *x*-coordinate so that *x*_1_ ≤ *x*_2_ ≤ ⋯ ≤ *x_N_*.

The best piecewise linear fit for the first *n* datapoints with *I* pieces corresponds to: (1) the best piecewise linear fit for the first *n*′ datapoints with *ℓ* – 1 pieces and (2) the best linear function fit for the remaining *n* – *n*′ datapoints, for some *n*′ < *n*. This yields the following recurrence:

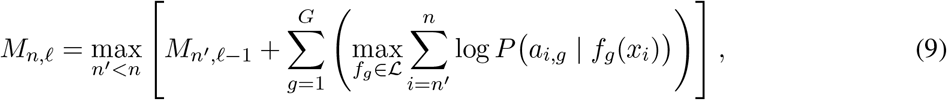

where the inner maximization is solved using regression.

The DP algorithm consists of using the recursion (9) to fill in a DP table column-by-column followed by a pass backwards through the table to identify the *L*-piecewise linear expression functions *f*_1_,…, *f_G_* and breakpoints *b*_1_,…, *b*_*L*–1_. The run-time of the DP algorithm is upper-bounded as *O*(*LN*^2^*G*·*P*_0_), where *P*_0_ is the runtime to solve the inner maximization in (9) for a single gene *g*. For the case where the gene expression values *a_i,g_* follow the Gaussian model, the run-time can be shortened to *O*(*LN*^2^*G*) by using linear algebra techniques from [2]. When *N* is large, it is also possible to reduce the run-time of the dynamic programming algorithm – at the expense of spatial resolution – by restricting the coordinates *n*′, *n* in the recurrence (9) to a subsequence of the *N* data points. In practice, in our analyses, we observe marginal loss in spatial resolution when restricting to subsequences of size 150 or larger.

## 6 Modeling *L*-layered tissues with conformal maps

For an arbitrary *L*-layered tissue slice with layer boundaries Γ_1_,… Γ_*L*–1_, we similarly assume that the expression of a gene *g* depends only on the layer depth, i.e., the distance to the layer boundaries Γ_*ℓ*_. However, because the layer boundaries Γ_*ℓ*_ are not necessarily axis-aligned, the depth cannot be immediately computed. An elegant solution is obtained by using a conformal map, a tool from complex analysis [4]. A conformal map 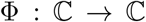 is a complex function that locally preserves angles; equivalently, Φ is conformal if it is analytic and has non-zero derivative everywhere. Note that the inverse Φ^−1^ of a conformal map Φ is also conformal. Conformal maps are used to solve differential equations in the plane 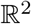 with complicated boundary conditions – e.g. heat flow in a plate or airflow around a wing – by identifying 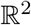 with the complex plane 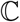 [68].

Here, we use conformal maps to transform a layered tissue slice *T* into an axis-aligned layered tissue slice *T*′. Ideally, we would derive a conformal map Φ : *T* → *T*′ that transforms tissue slice *T* to axis-aligned tissue slice *T*′, with the constraints that Φ maps each layer boundary Γ_*ℓ*_ in *T* to the corresponding layer boundary *x* = *b_ℓ_* in axis-aligned tissue slice *T*′. Unfortunately, constructing a conformal map satisfying these additional contraints is challenging, as the standard constructions of conformal maps only allow constraints on the boundary of *T* [68]. In general, it is unclear whether a conformal map satisfying additional constraints on non-boundary curves always exists.

As an alternative, we derive *L* conformal maps 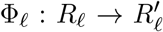 for *ℓ* = 1,…, *L*, where Φ_*ℓ*_ transforms layer *R_ℓ_* of tissue slice *T* to the corresponding layer 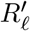 of axis-aligned tissue slice *T*′. We require that each conformal map Φ_*ℓ*_ maps the layer boundaries Γ_*ℓ*–1_, Γ_*ℓ*_ of layer *R_ℓ_* to the corresponding layer boundaries *x* = *b*_*ℓ*–1_, *x* = *b_ℓ_* of layer 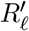, respectively, which is equivalent to the following constraints on the conformal map Φ*ℓ*:

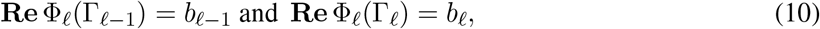

where **Re** denotes the real part of a complex number. We define 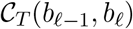 as the set of conformal maps Φ_*ℓ*_ that satisfy (10). Notice that (10) are constraints on the boundary of each layer *R_ℓ_*, and thus such maps are guaranteed to exist. Then, the map Φ : *T* → *T*′ from tissue slice *T* to axis-aligned tissue slice *T*′ is given by the piecewise sum of the *L* conformal maps Φ_1_,…, Φ_*L*_, that is

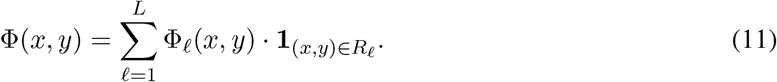

By the symmetry principle [68], if the limits of Φ_*ℓ*_ and Φ_*ℓ*+1_ at Γ_*ℓ*_ are the same and are continuous for all *ℓ*, then Φ is a conformal map. However, it is unknown whether these equality and continuity conditions are guaranteed to hold for arbitrary layer boundaries Γ_1_*,…*,Γ_*L*_.

Analogous to the axis-aligned setting, we call the real part **Re** Φ_*ℓ*_(*x, y*) of the conformal map Φ_*ℓ*_(*x, y*) the *relative* depth of position (*x, y*) in layer *R_ℓ_*. We model the expression of gene *g* in axis-aligned tissue slice *T*′ using a piecewise linear function *f_g_*(*x*) as in Section 5. Then, our goal is to find conformal maps Φ_1_, …, Φ_*L*_ and piecewise linear functions *f_g_*(*x*) such that the measured gene expression **A** at spots **S** in the tissue slice *T* are best fit by piecewise linear functions *f_g_*(*x*) in the corresponding axis-aligned tissue *T*′. We define the *L*-Layered Problem as the following maximum likelihood estimation problem.

### *L*-Layered Problem

*Given SRT data* (**A, S**), *layered tissue slice* 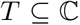, *and a number L of pieces, find an axis-aligned tissue slice T*′ *with layers x* = *b*_1_,…, *x* = *b*_*L*–1_, *conformal maps* 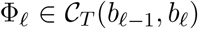, *and L-piecewise linear expression functions* 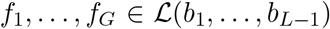 *that maximize the log-likelihood of the data*:

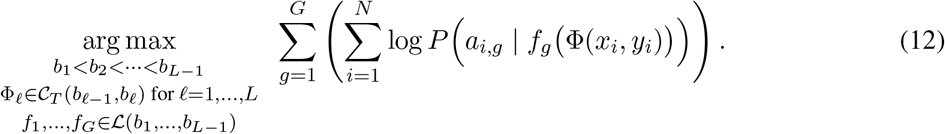

The *L*-Layered Problem generalizes the assumption in the Axis-Aligned *L*-Layered Problem that the expression function *f_g_*(*x,y*) of a gene *g* at position (*x,y*) depends only on the distance from the layer boundaries *x* = *b*_*ℓ*–1_ and *x* = *b_ℓ_*; in this more general formulation, the expression function *f_g_*(*x, y*) of gene *g* is constant along contours that “interpolate” between adjacent layer boundaries Γ_*ℓ*–1_ and Γ_*ℓ*_ (Figure 2B).

In general, solving the L-Layered Problem without additional constraints on the conformal maps Φ_1_,…, Φ_*L*_ is challenging. Below, we give algorithms for solving two special cases of *L*-Layered Problem which provide useful approximations on real data: (1) when the layer boundaries Γ_1_,…, Γ_L_–_1_ are known and (2) when the layer boundaries Γ_1_,…, Γ_*L*–1_ are lines.

## 6.1 Approximate layer boundaries Γ_*ℓ*_ are given

Suppose we are given prior information about the spatial organization of a tissue slice *T* in the form of approximate layer boundaries 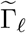. While these approximate layer boundaries may not directly correspond to the true layer boundaries Γ_*ℓ*_, they can nevertheless be used to compute conformal maps Φ_*ℓ*_. From these conformal maps, we can then estimate piecewise linear expression functions *f*_1_,…, *f_G_*, but without requiring that the breakpoints *b_ℓ_* of the piecewise linear functions *f_g_* occur at the mapped approximate layer boundaries Φ_*ℓ*_(Γ_*ℓ*_). In this way, we can incorporate prior information about tissue geometry (e.g. from prior knowledge of the tissue structure or from H&E images) without requiring that the expression functions conform exactly to this prior information.

Specifically, suppose we are given approximate layer boundaries 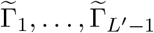, where *L*′ is not necessarily equal to the desired number *L* of layer boundaries. We constrain each conformal map Φ_*ℓ*_ so that the layer boundaries 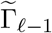 and 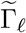 are mapped to lines 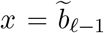 and 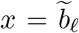, respectively, for some choice of real numbers 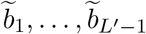. These constraints on the conformal maps Φ_*ℓ*_ are equivalent to the constraints 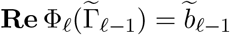 and 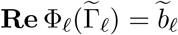 for all *ℓ* = 1,…, *L*′, which is in turn equivalent to the constraint 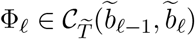, where 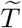 is tissue slice *T* equipped with the approximate layer boundaries 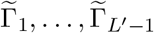 and approximate layers 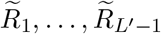. We define the conformal map 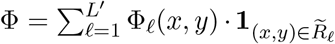 as the piecewise sum of the *L*′ conformal maps Φ_ℓ_.

Thus, given approximate layer boundaries 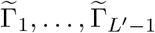 and lines 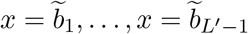, we solve the following optimization problem which we call the *L-Layered Problem with Approximate Layer Boundaries*:

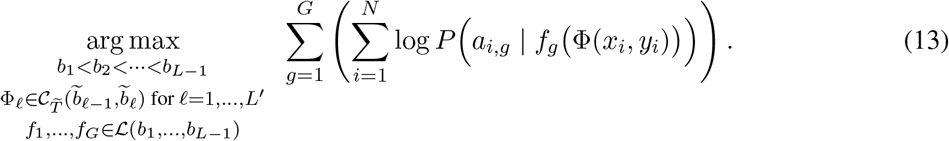

The conformal maps Φ_*ℓ*_ in (13) can be solved for separately from the breakpoints *b_ℓ_* and expression functions *f_g_* by using the observation that the constraint 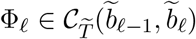 *uniquely* defines the real part of the conformal map Φ_*ℓ*_. This is because, in the constraints 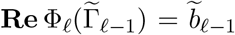 and 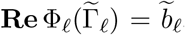, the real part **Re** Φ_*ℓ*_ of each conformal map Φ_*ℓ*_ is a harmonic function, and thus can be written as the solution to the heat equation with specified boundary conditions. That is, if one fixes curve 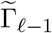 to have constant temperature 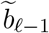 and 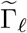 to have constant temperature 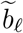, then **Re** Φ_*ℓ*_(*x, y*) is the temperature at point (*x, y*) and has a unique solution (Figure 2B). In practice, because there is no closed-form solution to the continuous heat equation for arbitrarily shaped boundaries, we solve the heat equation on a discretized tissue slice using a random walk-based approach [50, 36]. See the Appendix for more details.

After computing the conformal maps Φ_1_,…, Φ_*L*_′, the optimization problem in (13) reduces to

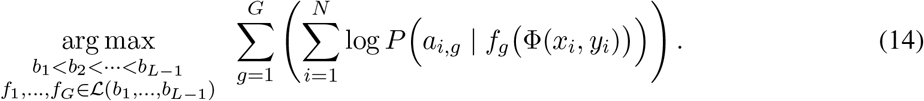

This is an instance of the Axis-Aligned *L*-Layered Problem with transcript counts *a_i,g_* and transformed spots Φ(**s**_*i*_). Thus, we solve (14) by using the dynamic programming algorithm for the Axis-Aligned *L*-Layered Problem (Section 5.2).

We note that in general, there are multiple ways to specify lines 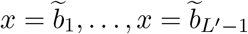 in the *L*-Layered Problem with Approximate Layer Boundaries. Two reasonable choices include: (1) defining *b*_1_,…, *b*_*L*′–1_ such that the difference 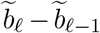 between consecutive lines is proportional to the physical distance between layer boundaries Γ_*ℓ*_and Γ_*ℓ*–1_, or (2) setting *b_ℓ_* = *ℓ* for *ℓ* = 1,…, *L*′. We note that while the optimization objective (13) is invariant under multiplicative scaling of the lines *b_ℓ_*, the estimated slopes 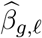 will differ. In the first choice, the estimated layer-specific slopes Γ_*g,ℓ*_ correspond to physical gradients of expression and are comparable both across genes g within each layer and across layers, while in the second choice the slopes are only comparable across genes *g* in the same layer *R_ℓ_*. In this work, we follow (1) and set 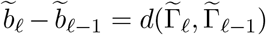 where 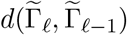 is the partial Hausdorff distance, a standard distance measure in computer vision [41].

## 6.2 Layer boundaries Γ_*ℓ*_ are lines

The second special case of the *L*-Layered Problem is when the layer boundaries Γ_1_,…, Γ_*L*–1_ are lines. Let 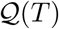 be the set of lines Γ with endpoints on the boundary *∂T* of the tissue *T*. In this case, given lines *x* = *b*_1_,…, *x* = *b*_*L*–1_, the *L*-Layered Problem reduces to solving the following optimization problem which we call the *Linear L-Layered Problem*:

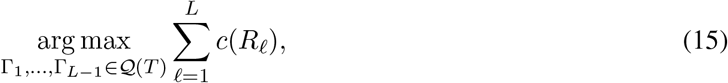

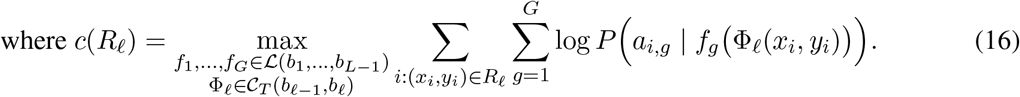

We derive a dynamic programming algorithm to solve (15) for any convex tissue slice *T* and any function *c*(*R*) that maps subsets *R* ⊆ *T* of the tissue to 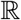. We note that the specific function *c*(*R*) in (16) can be computed by solving the heat equation as described in Section 6.1. Our dynamic programming algorithm generalizes the dynamic programming algorithm for the Axis-Aligned *L*-Layered Problem from Section 5.2.

Briefly, the dynamic programming algorithm for solving (15) is as follows. Without loss of generality, let **s**_1_,…, **s**_N_boundary__ be the spots on the boundary *∂T* of the tissue *T* in clockwise order. For convenience, we define [*n, m*] to be the sequence (*n* + 1,…, *m* – 1) of indices when *m* > *n*, and the sequence (*n* + 1,…, *N*_boundary_, 1,…, *m* – 1) when *n* > *m*. We define *T_n,m_* ⊆ *T* to be the region of the tissue *T* that is formed by drawing a line Γ between spots **s**_n_ and **s**_m_ and has boundary spots {**s**_*i*}*i*∈[*n,m*_].

Let *M_n,m,ℓ_* be the best fit with *ℓ* nested layers (i.e. with *ℓ* –1 linear layer boundaries) in the region *T_n,m_*, that is

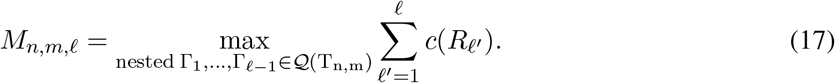

Then, the best fit with *ℓ* layers in the region *T_n,m_* can be decomposed to the sum of: (1) the best fit with *ℓ* – 1 layers in the region *T*_*n*′,*m*_′ and (2) the fit *c*(*T_n,m_* \ *T*_*n*′, *m*_′) for the region *T_n,m_* \ *T*_*n*′,*m*_′, for some *n*′, *m*′ ∈ [*n*, *m*]. Thus, we have the following recurrence relationship:

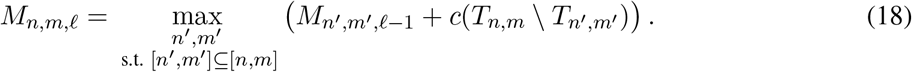

After computing the values *M_n,m,ℓ_* for all 1 ≤ *n, m* ≤ *N*_boundary_ and *ℓ* = 1,…, *L* – 1, the optimal value of (15) is then derived as

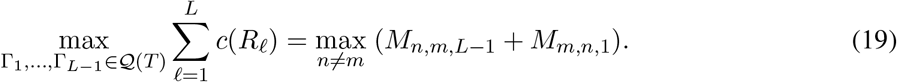

The dynamic programming algorithm consists of using the recursion (18) to fill in a table, followed by computing (19) to identify the best fit with *L* – 1 linear layer boundaries for the entire tissue *T*. See Figure S4 and the Appendix for more details.

### 6.2.1 Layer boundaries Γ_*ℓ*_ are parallel lines

We also derive a more computationally efficient algorithm for the case when the layer boundaries Γ_1_,…, Γ_*L*–1_ are *parallel* lines. In this case, each conformal map Φ_*ℓ*_ is a rotation map 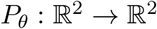 with the same angle *θ*, where *P_θ_*(*x, y*) rotates points (*x, y*) about the origin by angle *θ* ∈ [0, *π*). Then, the *L*-Layered Problem reduces to the following optimization problem which we call the *θ-Rotated L-Layered Problem*:

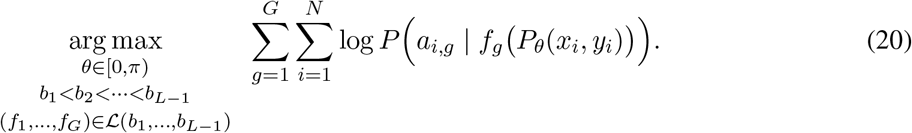

For a fixed angle *θ*, the optimization problem in (20) reduces to the Axis-Aligned *L*-Layered Problem with expression matrix *A* and rotated spots *Pθ(x,¿, yfi.* Thus, we solve the θ-Rotated L-Layered Problem by performing a parameter sweep over angles *θ ∈* [0, *π)* and finding the value of *θ* that maximizes the objective of (20).

## 7 Improving run-time with dimensionality reduction

The algorithms described above for solving different versions of the *L*-Layered Problem solve a separate regression problem for each gene. This can result in large run-times when the number of genes *G* is large, particularly when using a Poisson model for the UMI counts *a_i,g_*. This is because Poisson regression does not have a closed-form solution and requires solving a convex optimization problem, which is generally slower to solve than linear regression.

To reduce the run-time for SRT data (**A**, **S**), we use generalized principal components analysis (GLM-PCA) [87] to produce a low-dimensional representation **U** of the expression matrix **A**, and then run the algorithms presented above on the top-2L generalized principal components 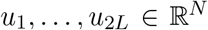 using the Gaussian model. This results in large reductions in run-time: for example, for the Axis-Aligned *L*-Layered Problem, we reduce the run-time of our DP algorithm from *O*(*LN*^2^*G* · *P*_0_) under the Poisson model to *O*(*LN*^2^ · 2*L*) = *O*(*N*^2^*L*^2^), corresponding to the run-time for DP with the Gaussian model and *G* = 2*L* genes. After computing the breakpoints *b*_1_,…, *b_L_*, we then estimate the expression functions *f_g_* for each gene *g* by computing *L* separate Poisson regressions as described in Section 5.1, for which the run-time is only *O*(*LGP*_0_). In practice, on the DLPFC Donor I data we observe a 100 ×reduction in run-time when solving the Axis-Aligned *L*-Layered Problem with GLM-PCA-reduced data matrix **U**, as opposed to the original data matrix **A**, with nearly identical performance in layer identification.

Our use of GLM-PCA is motivated by the observation that for SRT data (**A**, **S**) following the Poisson model with piecewise linear expression functions *f_g_* and breakpoints *b*_1_,…, *b*_*L*–1_, the top-2L generalized principal components *u*_1_,…, *u*_2*L*_ of the expression matrix **A** are approximately piecewise linear with breakpoints *b*_1_,…, *b*_*L*–1_. See the Appendix for more details.

## 8 Simulation details

For the first simulation, we simulated SRT data (**A**, **S**) from a rectangular tissue slice *T* with parallel layer boundaries lines Γ_1_, Γ_*L*–1_ that are rotated by an angle *θ* = 40° from the *x*-axis. The observed spots **S** are equally spaced in the tissue slice *T* and form a 50 × 40 grid. We generated piecewise linear expression functions *f_g_*(*x, y*) for *G* = 1000 genes with slopes *β_g,ℓ_* and *y*-intercepts *α_g,ℓ_* chosen uniformly at random from [−0.1,0.1]. We generated transcript counts *a_i,g_* using the Poisson model with added Gaussian noise 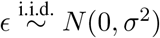

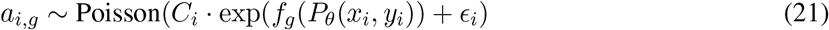

where *C_i_* was chosen so that each spot had total UMI count of approximately 2000. We ran Belayer to solve the *θ*-Rotated *L*-Layered Problem (Section 6.2.1), and compared Belayer to three other approaches: BayesSpace [99], SpaGCN [40], and the SCANPY [92] implementation of leiden clustering algorithm [88]. (We do not compare against stLearn [74] as an H&E image provides crucial information in its spatial cluster step, which smooths gene expression across space by weighted averaging where the weights are derived from H&E image.) We assume all methods know the true number *L* of layers. We assessed the performance of each method by computing the Adjusted Rand Index (ARI) between the *L* layers/clusters estimated by each method and the true layers.

For the second simulation, we evaluated Belayer on simulated SRT data (**A**, **S**) from a rectangular tissue slice *T* similar to the tissue slice described above, except *θ* is uniformly randomly chosen from 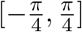, and the spots form a 70 × 50 grid. We generated transcript counts *a_i,g_* using Splatter [96] with *G* = 1000 genes such that: the expression function *f_g_*(*x, y*) is approximately piecewise linear in each layer for each gene *g* (using the trajectory simulation feature of Splatter and placing cells in each spot by the pseudotime of each trajectory), the median UMI count per spot is around 1100, and the probability that a gene has a different linear expression function across layers, which we call DE probability, is set to a fixed value using the differential expression probability parameter in Splatter. We generated 5 simulated SRT datasets (**A**, **S**) for each value of *L* = 4…, 8 and each value of DE probability in {0.1, 0,2, 0.3, 0.4, 0.5}. The details for running Belayer and comparing to other methods are the same as the first simulation.

## 9 Data processing and parameter selection

For Belayer, we choose the number L of layers by identifying the elbow in the consecutive differences of the negative log-likelihood. See Figure S7 for an example for DLPFC sample 151508 and the skin wound dataset.

We use the following data processing and parameter settings for running other SRT methods.

### BayesSpace

We use function `spatialPreprocess` implemented in BayesSpace for data pre-processing. This function normalizes the count matrix, selects a given number of highly variable genes, and computes a given number of principle components using PCA. We compute BayesSpace clustering results with both 2000 highly variable genes for real SRT datasets, and use all genes for simulated data. We follow the instructions in BayesSpace paper and set the number of PCs to be 15 for DLPFC dataset, and we use 2*L* PCs for simulated data and other SRT datasets. The number of clusters is either set to be the same as *L* chosen by Belayer or selected by its model selection function qTune in real SRT datasets, and set to be the true number of clusters in simulated data.

### SCANPY

We follow the SCANPY tutorial to pre-process DLPFC and simulated data. The UMI count matrix is normalized to a target sum of 10^6^ per spot for the DLPFC data and 10^3^ per spot for the simulated data, and then transformed by log1p. We use all genes to compute PCs using SCANPY with the default number of PCs. We then use SCANPY to construct a neighborhood graph with various number of neighbors, and apply leiden algorithm in SCANPY to the neighborhood graph for clustering with various resolution parameter. Since SCANPY does not have its own model selection function, we vary the number of neighbors parameter within the range of 10 and 50 and vary the resolution parameter within the range of 0.1 and 1.5 until the number of clusters matches the number *L* of layers chosen by Belayer.

### stLearn

We follow the stSME clustering tutorial of stLearn to cluster DLPFC, mouse somatosensory, and mouse skin data. Briefly, stSME clustering of stLearn first combines spatial location and morphology from image to normalize gene expression, and then applies PCA and KMeans clustering to the normalized gene expression. Slide-SeqV2 does not generate H&E images, and stLearn uses only spatial location for gene expression normalization and further clustering.

### SpaGCN

We follow the SpaGCN tutorial for applying SpaGCN to 10X Visium and Slide-Seq data. As suggested by the SpaGCN tutorial and manuscript, we use parameter *p* = 0.5 for 10X Visium data and *p* = 1 for Slide-SeqV2 data.

### HotSpot

We run HotSpot with the default settings (as suggested by the Github repository): setting the null model to be the depth-adjusted negative binomial model and setting the number of neighbors in the *k*-NN graph to be 30. We rank genes by their estimated FDRs.

### SpatialDE

We follow the instructions of SpatialDE github for normalizing SRT data and running SpatialDE. We use the variance stabilizing normalization implemented in `NaiveDE.stabilize` for normalizing UMI count matrix, and then regress out the log of total UMI count per spot by `NaiveDE.regress_out`. SpatialDE is then applied to compute the p-value and q-value for each gene being a spatially varying gene.

### SPARK

SPARK uses a Poisson distribution to model the count data, and thus normalization is not needed. We apply SPARK to the UMI count data directly, and set the gene-filtering parameter in SPARK to filter out genes that are expressed in less than 10% of spots or have less than 10 UMI counts in total.

## 10 Code and Data Availability

Belayer is available at github.com/raphael-group/belayer. The DLPFC dataset is available at http://spatial.libd.org/spatialLIBD/. The mouse skin wound dataset can be accessed from GEO with accession GSE178758. The full size Slide-SeqV2 mouse cortex dataset is available at https://singlecell.broadinstitute.org/single_cell/study/SCP815/highly-sensitive-spatial-transcriptomics-at-near-cellular-resolution-with-slide-seqv2, and the code to extract the mouse somatosensory cortex is located ast https://github.com/dmcable/spacexr/blob/master/AnalysisPaper/SuppFigures/supp_part2.Rmd in the section “Somatosensory Cortex Cortical Cell Types”.

## 11 Acknowledgements

U.C. is supported by NSF GRFP DGE 2039656. This research is supported by National Cancer Institute (NCI) grant U24CA264027 to B.J.R.

## Supplemental Information

### A Relationship to Gaussian Process Models

Piecewise linear functions can be modeled using a Gaussian process (GP) having a one-dimensional linear covariance function *k*(**s**,**s**′) of the form

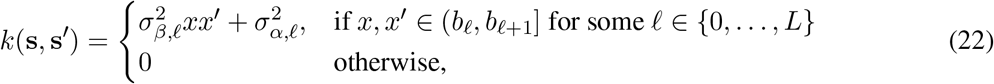

for breakpoints ∞ = *b*_0_ < *b*_1_ < ⋯ *b*_*L*–1_ < *b_L_* = ∞, where *x* and *x*′ are the *x*-coordinates of spots **s** = (*x,y*) and **s** = (*x*′, *y*′), respectively. Note that the linear covariance function is different from the widely used squared exponential covariance function, which is defined as 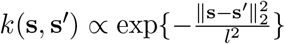.

Sampling an expression function *f_g_*(*x*) from the distribution 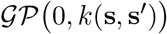 with covariance function (22) generates a piecewise linear model of the form (4) with Gaussian priors 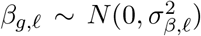 and 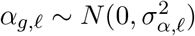 on the slope *β_g,ℓ_* and *y*-intercept *α_g,ℓ_*, respectively, of the gene expression in layer *ℓ*.

Using Gaussian Process (GP) prior 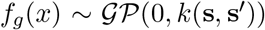, we define the following maximum a posteriori (MAP) version of the Axis-Aligned *L*-Layered Problem.

#### Axis-Aligned *L*-Layered Problem (MAP Version)

*Given SRT data* (**A**,**S**) *and a number L of layers, find layer boundaries x* = *b*_1_,…, *x* = *b*_*L*–1_ *and piecewise linear expression functions* 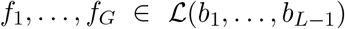 *that maximize the posterior distribution of the data under the prior* 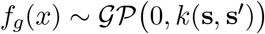 *for covariance function k*(**s**,**s**′) *in* (22):

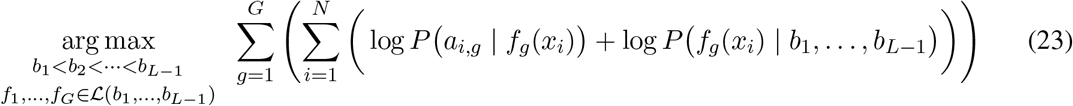

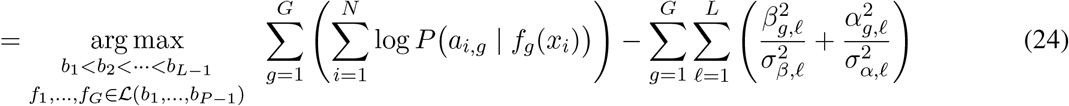

### B Dimensionality reduction using GLM-PCA

Let (**A**, **S**) be SRT data generated from an axis-aligned tissue slice *T* with breakpoints *b*_1_,…, *b*_*L*–1_ and *L*-piecewise expression functions *f_g_*(*x*). Define the *mean expression matrix* 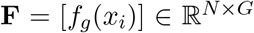 as the matrix of expression functions *f_g_*(*x*) evaluated at spatial locations *x* = *x_i_*. Given integer *K* > 0, GLM-PCA aims to find matrices **U**, **V** such that **F** ≈ **UV**, i.e. **UV** is a rank-*K* approximation of the mean expression matrix **F**. The columns *u_i_* of **U** are the *generalized principal components* of **A**.

In the Proposition below, we show that **F** has rank 2*L* by constructing an explicit rank decomposition **F** = **UV** where the columns *u_i_* of **U** are *piecewise linear*. We note that our rank decomposition in Proposition 2 does not correspond to the matrices **U**, **V** found by GLM-PCA; nevertheless, we empirically observe that the top-2*L* generalized principal components of **A**are approximately piecewise linear (Figure S1).

#### Proposition 2.

*Let* (**A**, **S**) *be SRT data generated from an axis-aligned L-layered tissue slice T with layer boundaries *x* = *b*_1_,…, *x* = *b*_*L*–1_ and expression functions *f_g_*(*x*). Let* 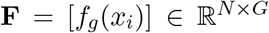 *be the mean expression matrix. Then* **F** *has a decomposition of the form* **F** = **UV** *for matrices* 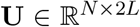 and 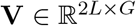, *where the columns* 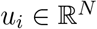 *of* **U** *are piecewise linear with breakpoints b_1_, …, *b*_*L*–1_.*

*Proof.* Without loss of generality assume that the spots are ordered by increasing *x*-coordinate, i.e. *x*_1_ ≤ *x*_2_ ≤ ⋯ *x_N_*. We define vectors 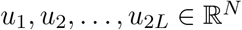 such that

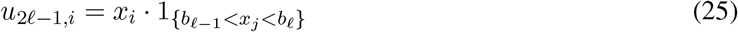

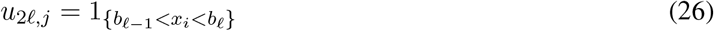

for all *ℓ* = 1,…, *L* and *i* = 1,…, *N* We similarly define vectors 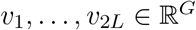 as

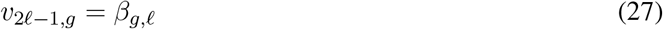

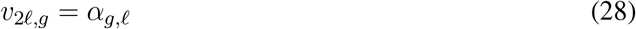

for all *ℓ* = 1,…, *L* and *g* = 1,…, *G*.

We define matrices 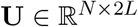 and 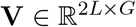 with columns 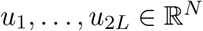 and rows 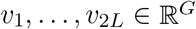, respectively. Then

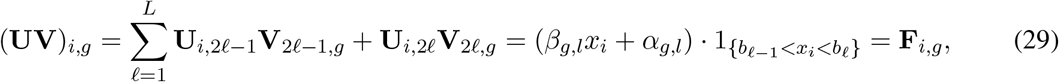

as desired.

**Figure S1:**
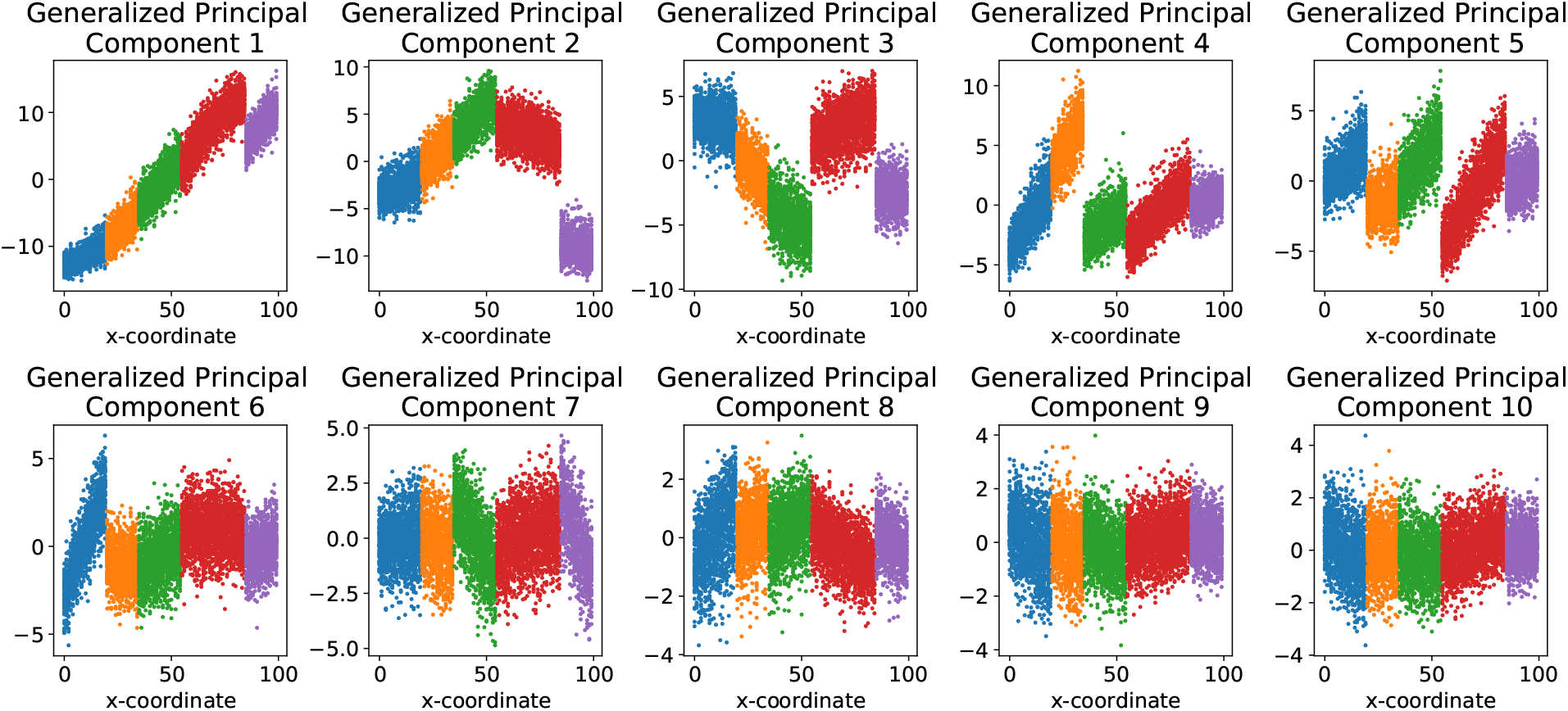
Top 2*L* generalized principal components for simulated SRT data (**A**, **S**) generated from an axis-aligned *L*-layered tissue slice *T* with *L* = 5 layers.

### C Improving visualization by binning spots

In our visualizations, we bin the expression values *a_i,g_* of spots **s**_*i*_ = (*x_i_*, *s_y_*) with similar *x*-coordinates *x_i_*. Specifically, we divide the range [min_*i*_ *x_i_*, max_*i x_i_*_] of *x*-coordinates into *N*_binned_ equal-width bins, where *B_j_* ⊄ [*N*] is the set of spots **s**_*i*_ = (*x_i_*, *y_i_*) in bin *j* and 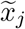 is the center of bin *j*. We then construct “binned expressions” 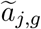 for each gene *g* and bin *j*, where 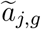 is the maximum likelihood estimate of the expression at *x*-coordinate 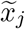, given the expressions {*a_i,g_*_}*i*∈*B_j_*_ of spots in bin *j* and under the assumption these spots have *x*-coordinate equal to 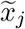. If the expressions *a_i,g_* follow the Poisson expression model, then the binned expression 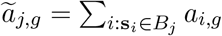 is the total expression of all spots in bin *j*, while if the expressions follow the Gaussian expression model, then the binned expression 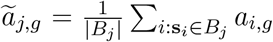 is the average expression of all spots in bin *j* (see e.g. Chapter 36 of [16]).

The binned expressions 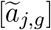 follow the same expression model and have approximately the same expression functions as the original expressions [*a_i,g_*], with the expression functions being equal for a sufficiently large number *N*_binned_ of bins [16]. Moreover, it is easier to visualize the piecewise linear expression functions *f_g_*(*x*) in the binned expressions 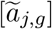 versus the unbinned expressions [*a_i,g_*] (Figure S2).

**Figure S2:**
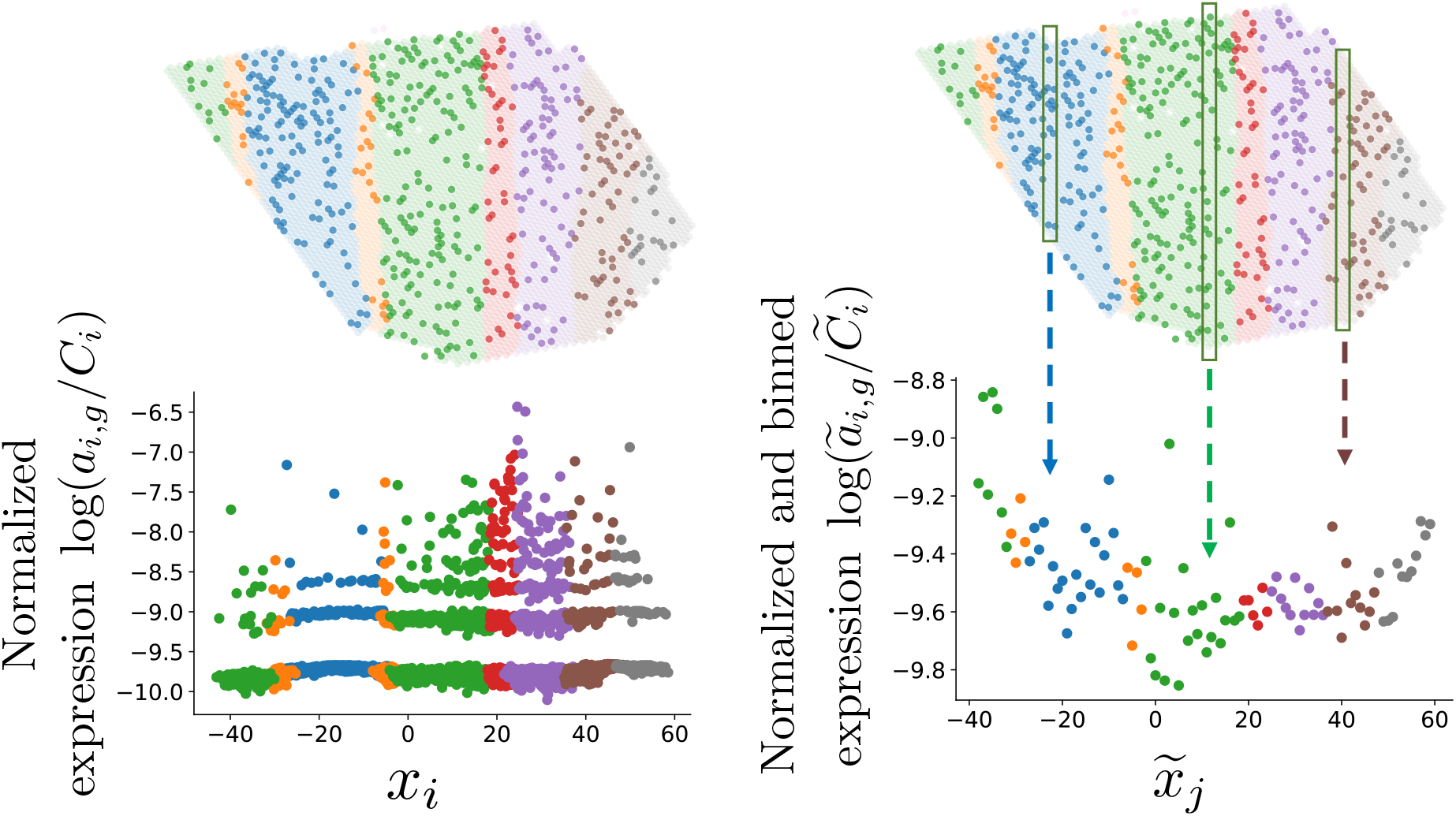
(Left) Visualization of *x*-coordinate versus normalized expression values from an axis-aligned layered tissue slice. (Right) Left plot with expressions binned across spots (*x, y*) with similar *x*-coordinates.

### D Solving the *L*-Layered Problem with Approximate Layer Boundaries

#### D.1 Solving heat equation on a discretized tissue slices with boundary conditions

As described in Section 6.1, we estimate the real part *u* = **Re** Φ_*ℓ*_ of each conformal map Φ_*ℓ*_(*x, y*) = *u* + *iv* by solving the following heat equation:

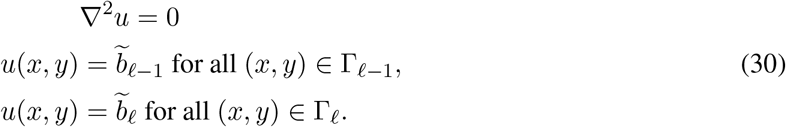

We solve (30) using the finite difference method [84], which estimates the values *u*(*x, y*) at grid points on either a regular square grid [84] or a regular hexagonal grid [52]. For our analyses, the two datasets obtained using the 10X Visium platform have spots that form a regular hexagonal grid. For the Slide-SeqV2 dataset, we overlay a regular square grid on the tissue and approximate the harmonic function value of each spot by that of its nearest grid point. We address the special cases of *ℓ* = 1, *L* at the end of this section.

**Figure S3:**
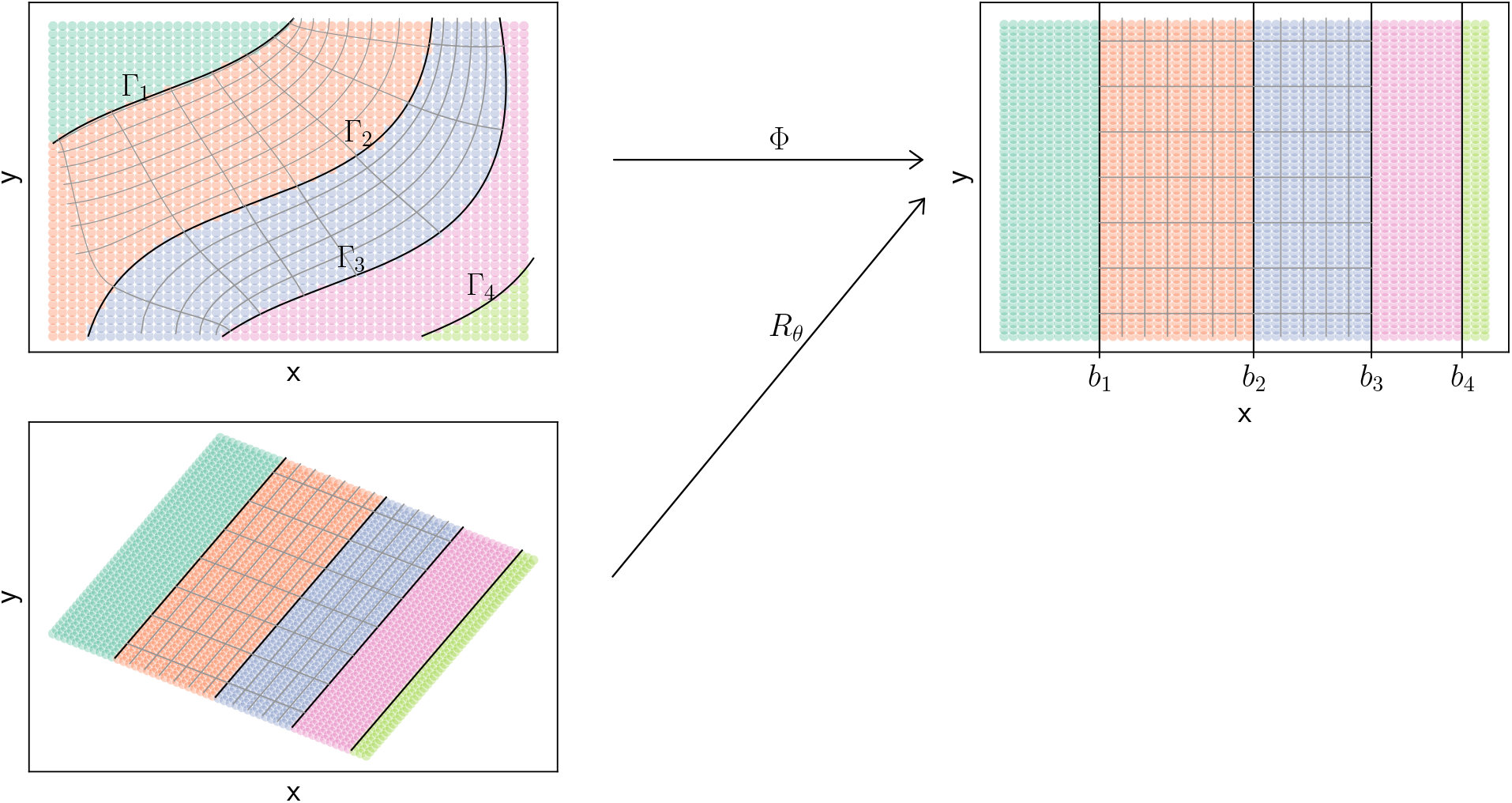
A conformal map Φ maps a 5-layered tissue slice to an axis-aligned tissue slice. Color indicates layers. The gray lines of the right panel in the second and third layer are grid lines parallel to *x* and *y* coordinate, their corresponding lines before mapping Φ are also known in gray lines. A special case when Φ is a rotation is shown in the lower panel.

We now briefly describe the finite difference method we implement to solve (30). Abusing notation, let *s*_1_, *s*_2_,…, *s_N_* be the grid points (from either a square grid or a hexagonal grid). Let *B*_*ℓ*–1_ be the grid points closest to/lying on the layer boundary Γ_*ℓ*–1_. We define *B_ℓ_* similarly. Let **u** = (*u*(*s*_1_),*u*(*s*_2_),…, *u*(*s_N_*)) be the vector of the function *u*(*x, y*) evaluated at grid points *s_i_*, let **u***B*_*ℓ*–1_, **u**_*B*_*ℓ*__ be the subset of **u** corresponding to *u*(*x, y*) evaluated at points in *B*_*ℓ*–1_, *B_ℓ_*, respectively. Let **u**_*I*_ be the function *u*(*x, y*) evaluated at points not in *B*_*ℓ*–1_, *B_ℓ_*, i.e. the interior points. Let *E_i_* be the set of grid points adjacent to *s_i_*. In a regular hexagonal grid case, |*E_i_*| = 6, while in a regular square grid case, |*E_i_*| = 4. We partition the vector **u** into two subsets: the spots **u**_*B*_

We discretize (30) into a system of linear equations of the form

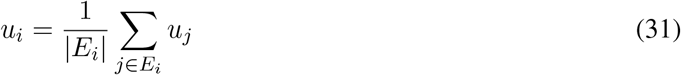

for all *i* = 1,…, *N*, where we constrain **u***B*_*ℓ*–1_, **u**_*B_ℓ_*_ as

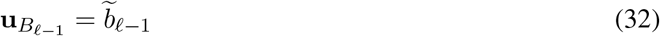

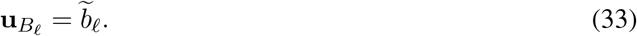

This linear system can be rewritten in matrix form as *L***u** = 0, where *L* is the Laplacian matrix of the grid, i.e.

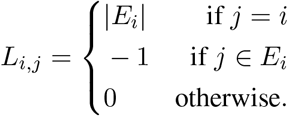

We also partition the Laplacian matrix *L* as 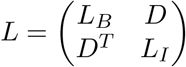, where *B* = *B*_*ℓ*–1_∪*B_ℓ_*. Then, the solution to the discretized heat equation (31) is given by

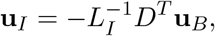

where **u**_B_ is constrained by (32), (33).

For the first layer *R*_1_ and the last layer *R_L_*, these regions are bounded by a layer boundary Γ_*ℓ*_ on one side and a tissue boundary *∂T* on all other sides. Moreover, the tissue boundaries are generated by tissue cutting and preparation in experiments and they may not be parallel to layer boundaries. Therefore, tissue boundaries are not informative of relative layer depth, and we do not solve the heat equation (30) with two boundaries condition We instead define the layer depth by heat diffusion from the one layer boundary, for which the temperature is fixed. As before, we denote the spots on the layer boundary by B and interior spots or the ones on tissue boundaries as *I*, and we partition the temperature **u** of spots by **u**_*B*_ and **u**_*I*_. Let *W* be the weighted adjacency matrix defined by

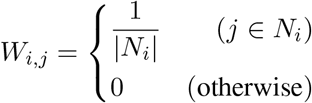

for *i* ∈ *I* and

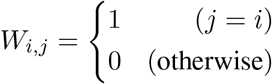

for *i* ∈ *B*. Without loss of generality, we assume that spots in *B* are placed on the top left of *W*. The temperature of spots in *I* after *t* step diffusion is 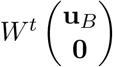. We use the diffusion after a fixed number of diffusion steps to define relative layer depth. The number of diffusion steps *t* is chosen as the smallest number of steps such that min **u**_*I*_ = *δ* min **u**_*B*_. That is, all spots have a nonzero temperature (δ is chosen to be 0.01) but far from reaching steady state. Note that the following regression is invariant by any affine transformation of **u**. We choose an affine transformation such that the largest difference of temperature max **u** – min **u** is the partial Hausdorff distance.

### E Dynamic programming algorithm for Linear *L*-Layered Problem

We derive a DP algorithm for solving an optimization of the form (15) for any function *c*(*R*) that maps subsets *R* ⊆ *T* of the tissue to 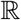. We present the algorithm for the case where the tissue slice *T* is a circle, but we note that the DP algorithm is directly applicable to any convex shape *T*.

#### Circle Problem Statement

We are given a circle *T* with *b* = *N*_boundary_ fixed points around its circumference *∂T* and an objective function *c*(*R*). We define 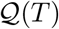 to be the set of lines Γ with endpoints on the boundary *∂T* of the tissue *T*. Without loss of generality, let the points on *∂T* be labeled clockwise **s**_1_,…, **s**_*b*_ in clockwise order. We expect 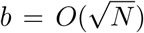, where *N* is the total number of points in the interior of *T*. Note that all arithmetic regarding points in *∂T* will be done mod *b*, unless otherwise stated. Let *M_L_* be the best fit with *L* layers (i.e., with *L* – 1 linear layer boundaries) in *T*. Then our goal is to find:

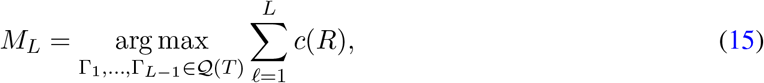

#### Circular Segment Dynamic Program

For convenience, we define [*n, m*] to be the sequence (*n* + 1,…, *m* – 1) of indices when *m* > *n*, and the sequence (*n* + 1,… , *N*_boundary_, 1,…, *m* – 1) when *n* > *m*. We define the circular segment *T_n,m_* ⊆ *T* to be the region of the tissue *T* that is formed by drawing a line Γ between spots **s**_*n*_ and **s**_*m*_ and has boundary spots {**s**_i_}_*i*_∈[_*n,m*_]. Call layer boundaries **Γ** = Γ_1_,… ,Γ_*ℓ*–1_ *nested* if for any two lines 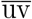 and 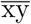 in **Γ** defined such that if *x* is the closest clockwise point to *u* in {*x, y, v*}, it is the case that *y* is the closest counterclockwise point to *v* in {*x, y, u*}. Note that nestedness is a stronger condition than the layer boundaries **Γ** not intersecting; see Figure S4A for an example of layer boundaries that are non-intersecting but are also not nested.

First, we consider the computation of the following related quantity (Figure S4B). Let *M_n,m,ℓ_* be the optimal nested layer boundaries with *ℓ* layers in the region *T_n,m_*, i.e.

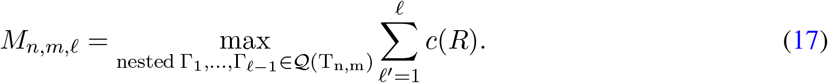

We first give a dynamic programming algorithm for computing *M_n,m,ℓ_* in *O*(*b*^4^ · *ℓ*) time. We will later leverage this algorithm in order to compute *M_L_*.

The best fit with *ℓ* layers in the region *Tℓ_n,m_* can be decomposed as the sum of: (1) the best fit with *ℓ* – 1 layers in the region *T*_*n*′, *m*′_ and (2) the fit *c*(*T_n,m_* \ *T*_*n*′, *m*′_) for the region *T_n,m_* \ *T*_*n*′, *m*′_, for some *n*′, *m*′ ∈ [*n, m*]. Thus, we have the following recurrence relationship:

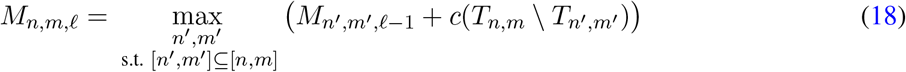

Our base cases are:

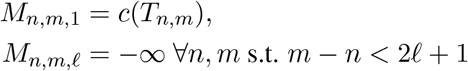

The first base case says if we have a circular segment *T_n,m_* with 1 layer (i.e. 0 layer boundaries), then that layer must include the entire circular segment. The second base case says that if there are not enough possible endpoints between *n* and *m* to place *ℓ* lines, then there is no possible solution. To see why, note that the endpoints of any layer boundary in *T_n,m_* must be in *V* = [*n* + 1, *m* – 1] and thus |*V*| = *m* – *n* – 1. For *M_n,m,ℓ_*, we must add *ℓ* lines using the endpoints in *V*, each with 2 endpoints. 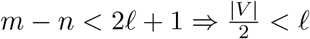, which implies there are not enough possible endpoints to add *ℓ* lines. By the pigeonhole principle, there is no possible solution for *M_n,m,ℓ_*.

Our recurrence (18) finds the *ℓ* best nested layer boundaries contained within *T_n,m_*. If the outermost layer boundary (i.e. the layer boundary closest to *n, m*) has endpoints *n*′, *m*′, then we must place an additional *ℓ* – 1 layer boundaries within *T*_*n*′, *m*′_, otherwise the layer boundaries would not be nested. The optimal value of the objective function in (18) is then the optimal value of placing *ℓ* ‒ 1 lines within *T*_*n*′*m*′_ added to the value of the objective function for the region *T_n,m_*\ *T*_*n*′ \, *m*′_. We check all possible values of *n*′, *m*′ and save the maximal value. See Figure S4B for a visualization of the recurrence (18).

**Figure S4:**
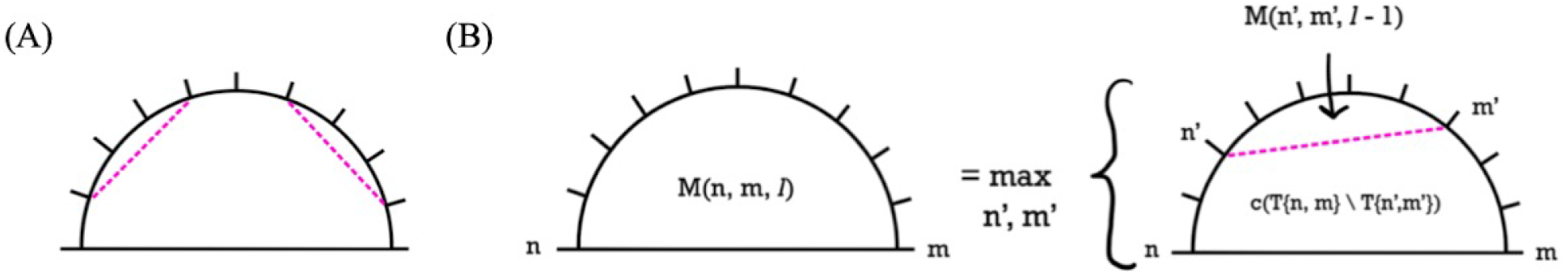

(**A**) Example of non-nested layer boundaries. (**B**) Visualization of recurrence for dynamic programming algorithm for linear layer boundaries.

#### Finding the First Layer Boundary

We find the optimal layer boundaries for the circle *T* by adding the value of the optimal nested layers for circular segment *T_n,m_* with the value of its complementary region *T_m,n_* for all *n, m* and taking the maximal result:

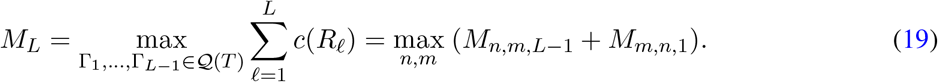

Note that the optimal set of layer boundaries has some layer boundary 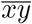 such that all other layer boundaries have endpoints in [*x, y*]. Because we are maximizing over all starting values of *n, m*, we are guaranteed to use *x, y* as the starting value at some point during the maximization of (19).

The above maximization (19) computes in *O*(*b*^2^) time, assuming a constant-time lookup for *M_n,m,ℓ_*. Furthermore, we construct a lookup table for all values of *M_n,m,ℓ_* for *n, m* = 1,…, *b* and *ℓ* = 1,…, *L* in time *O*(*b*^4^ · *L*) using the following algorithm:

#### Algorithm 1

Creating lookup table for all *M_n,m,ℓ_*

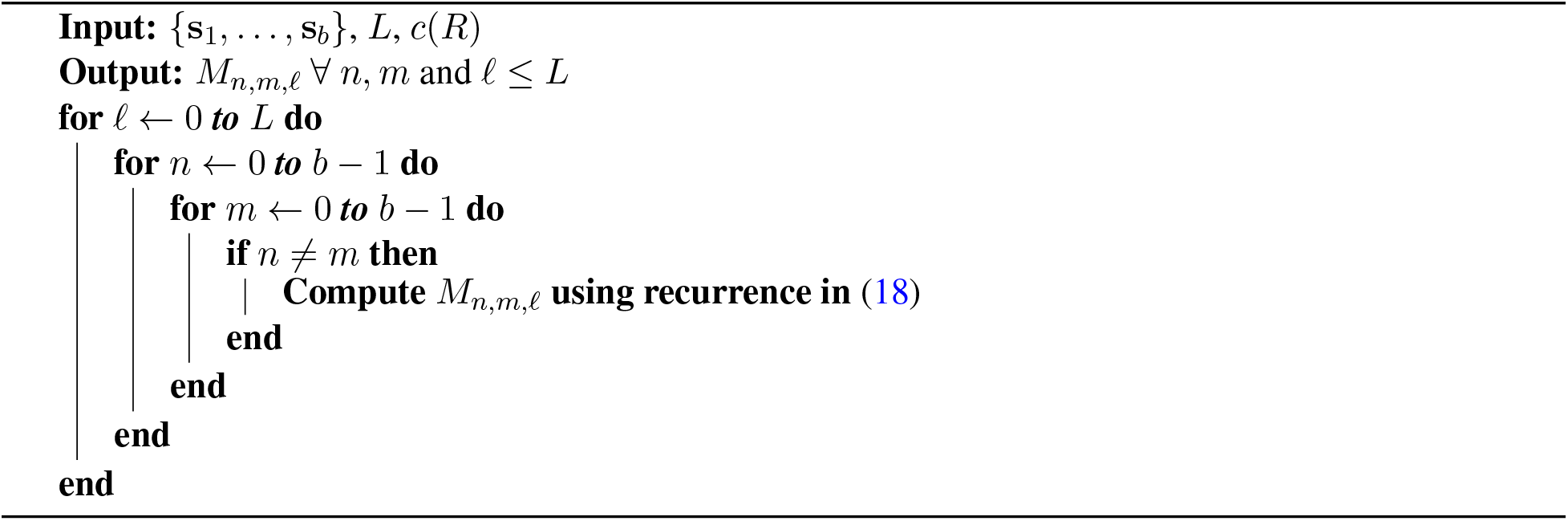

Note that for any *ℓ* ∈ {1,…, *L*}, the values of *M*_*n,m,ℓ*′_ for all *n, m*, and *ℓ*′ < *ℓ* will be populated in the lookup table before they are referenced by Algorithm 1 to compute *M_n,m,ℓ_*. In Algorithm 1, *n* and *m* each contribute a factor of *b* to the run-time. The inner maximization in the recurrence contributes a factor of *b*^2^, and iterating over *ℓ* = 1,…, *L* contributes a factor of *L*, so that Algorithm 1 has a run-time of *O*(*b*^4^ · *L*). Therefore, the overall run-time to find *M_L_* = *O*(*b*^4^ · *L*) + *O*(*b*^2^) = *O*(*b*^4^ · *L*). We note that this approach is easily adapted to any convex shape *T* by adding the constraint that no layer boundary can have endpoints on the same edge of the shape.

In practice, for the DLPFC analysis (Figure 3), computing *c*(*T_n,m_* \ *T*_*n*′, _*m*_′) takes approximately 20 hours on a 100-node cluster while computing *M_n,m,ℓ_*, and *M_L_* takes approximately 5*L* minutes in total. The bottleneck in computing *c*(*T_n,m_* \ *T_n_′ \ *m*′*_) is solving large linear systems in the heat equation. Improving the computation time is one future direction for improving Belayer.

### F Additional approaches for marker gene identification

We additionally evaluated (Figure S10) five other approaches for ranking genes in marker gene identification. We describe these ranking approaches below.

- Ranking genes by their Moran’s I score, a standard measure of spatial autocorrelation [67]
- Ranking genes by their Geary’s C score, another standard measure of spatial autocorrelation [35]
- Ranking genes by their likelihood ratio test (LRT) p-value, where we compare the maximum loglikelihood (12) with a piecewise constant expression function *f_g_* to the maximum log-likelihood (12) with a piecewise linear expression function *f_g_*.
- Ranking genes by their LRT p-value as above but restricted to Belayer Layer 3 (Figure 3D)
- Ranking genes by the sum 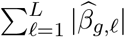 of their (absolute) layer-specific slopes across the *L* layers identified by Belayer.
- Ranking genes by the maximum difference 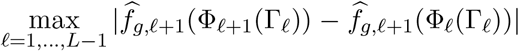 between layer-specific expression functions 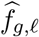) at layer boundaries Γ_*ℓ*_ across the *L* layers identified by Belayer.

**Figure S5:**
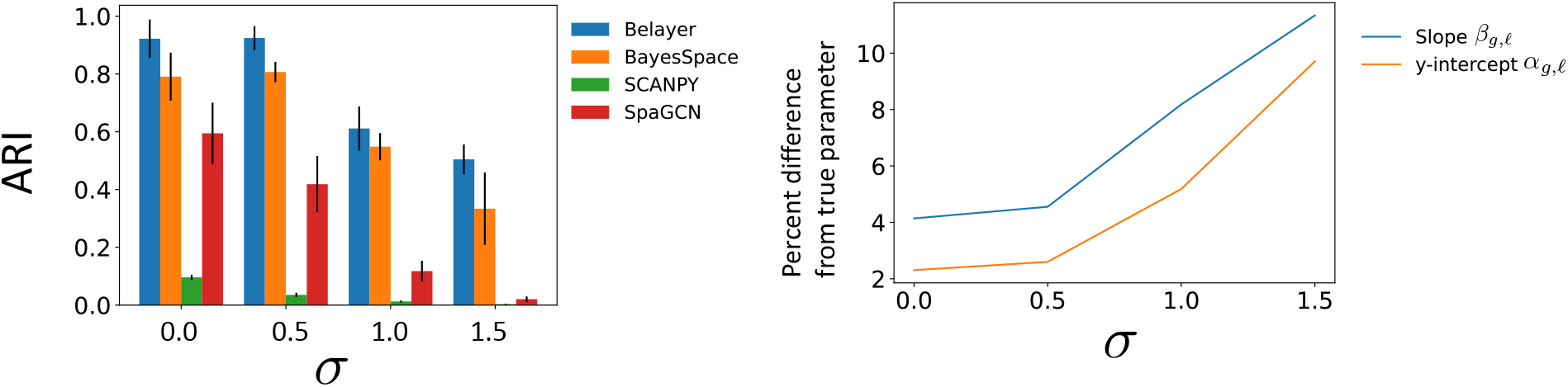
Comparison of Belayer, BayesSpace, SCANPY, and SpaGCN in identifying spatially distinct cell clusters in the first simulation. Performance of each method is evaluated according to the Adjusted Rand Index (ARI) and shown for different values of the number *L* of layers and standard deviation *σ* of the added Gaussian noise. Error bars indicate variation from 5 randomly simulated datasets for each parameter setting.

**Figure S6:**
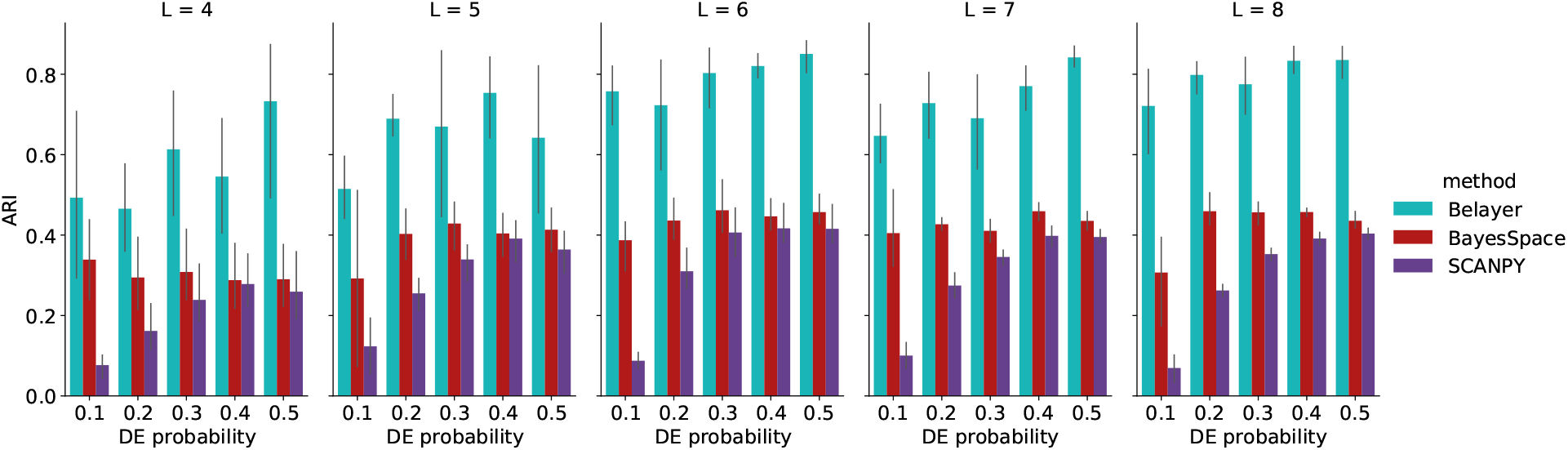
Comparison of Belayer, BayesSpace, and SCANPY in identifying spatially distinct cell clusters in the second simulation. Performance of each method is evaluated according to the Adjusted Rand Index (ARI) and shown for different values of the number *L* of layers and differential expression (DE) probability. Error bars indicate variation from 5 randomly simulated datasets for each parameter setting.

**Figure S7:**
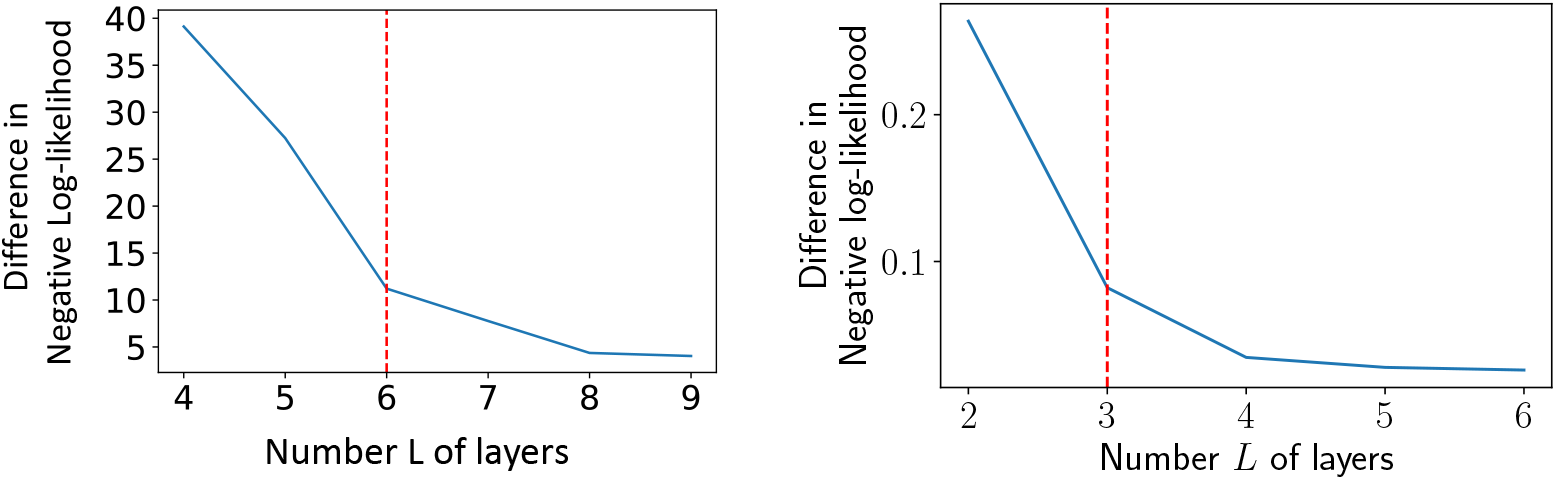
Number *L* of layers vs. difference in negative log-likelihood at *L* layers and *L* – 1 layers for DLPFC sample 151508 (left) and the mouse skin wound dataset (right). Vertical dashed line indicates manually annotated elbow.

**Figure S8:**
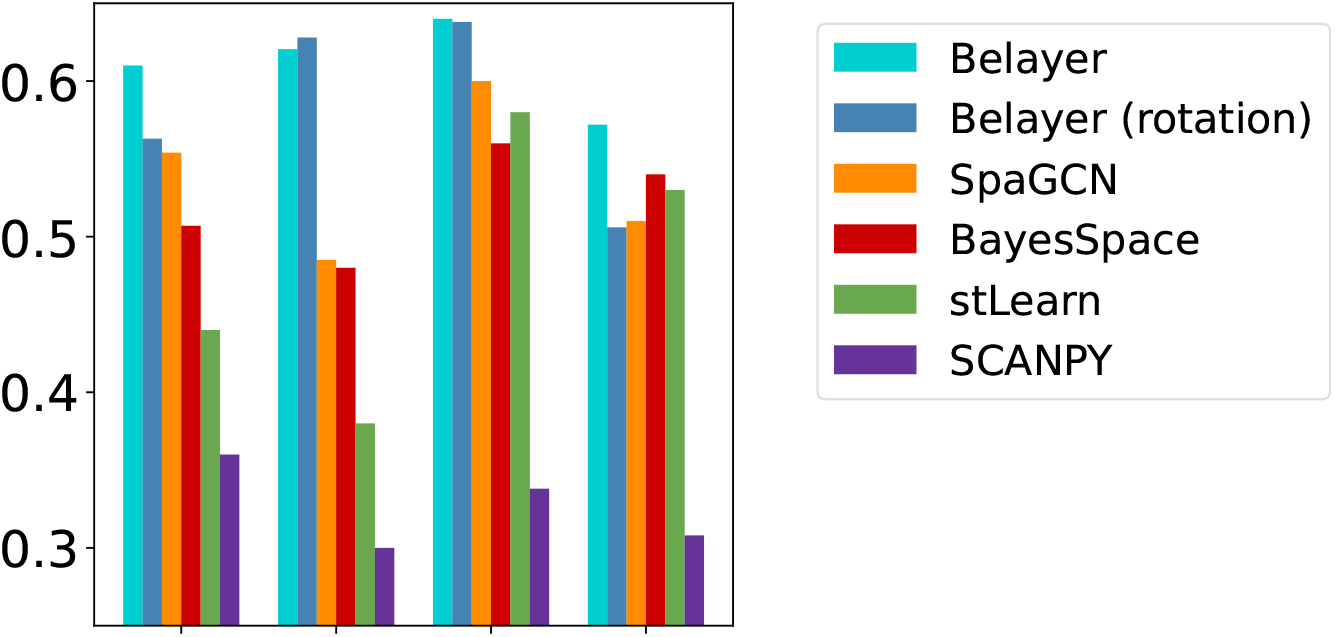
Comparison of Belayer (solving the Linear *L*-Layered Problem), Belayer (solving the θ-Rotated *L*-Layered Problem) and other methods in identifying annotated layers in SRT data from the DLPFC of Donor 1 as in Figure 3.

**Table S1:** Comparisons of Belayer, BayesSpace with Belayer’s model selection, and BayesSpace with its own model selection in identifying cortical layers of DLPFC data. Two samples in Donor 1 are excluded from this table because both model selections choose the same number of layers. The first two rows match the ARIs in Figure 3A.

|  | Donor 1 |  | Donor 2 |  |  |  | Donor 3 |  |  |  |
| --- | --- | --- | --- | --- | --- | --- | --- | --- | --- | --- |
|  | 151507 | 151508 | 151669 | 151670 | 151671 | 151672 | 151673 | 151674 | 151675 | 151676 |
| Belaye ARI<br>(Belaye model selection) | 0.61 | 0.621 | 0.356 | 0.61 | 0.707 | 0.596 | 0.653 | 0.669 | 0.697 | 0.551 |
| BayesSpace ARI<br>(Belaye model selection) | 0.507 | 0.48 | 0.322 | 0.363 | 0.403 | 0.505 | 0.364 | 0.338 | 0.329 | 0.459 |
| BayesSpace ARI<br>(BayesSpace model selection) | 0.482 | 0.472 | 0.257 | 0.230 | 0.398 | 0.484 | 0.528 | 0.322 | 0.425 | 0.406 |

**Figure S9:**
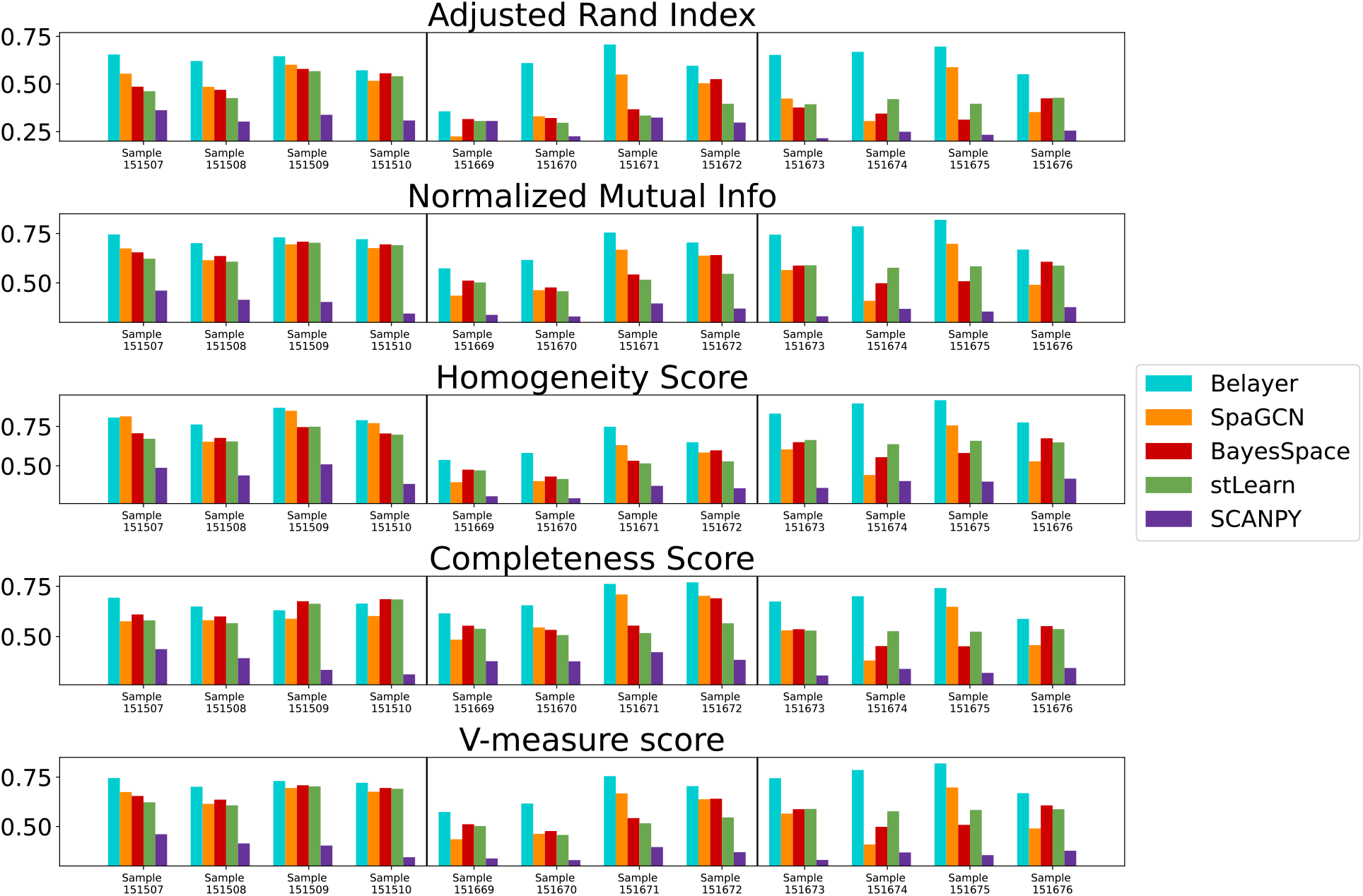
Comparison of Belayer and other methods from Figure 3 with different metrics.

**Figure S10:**
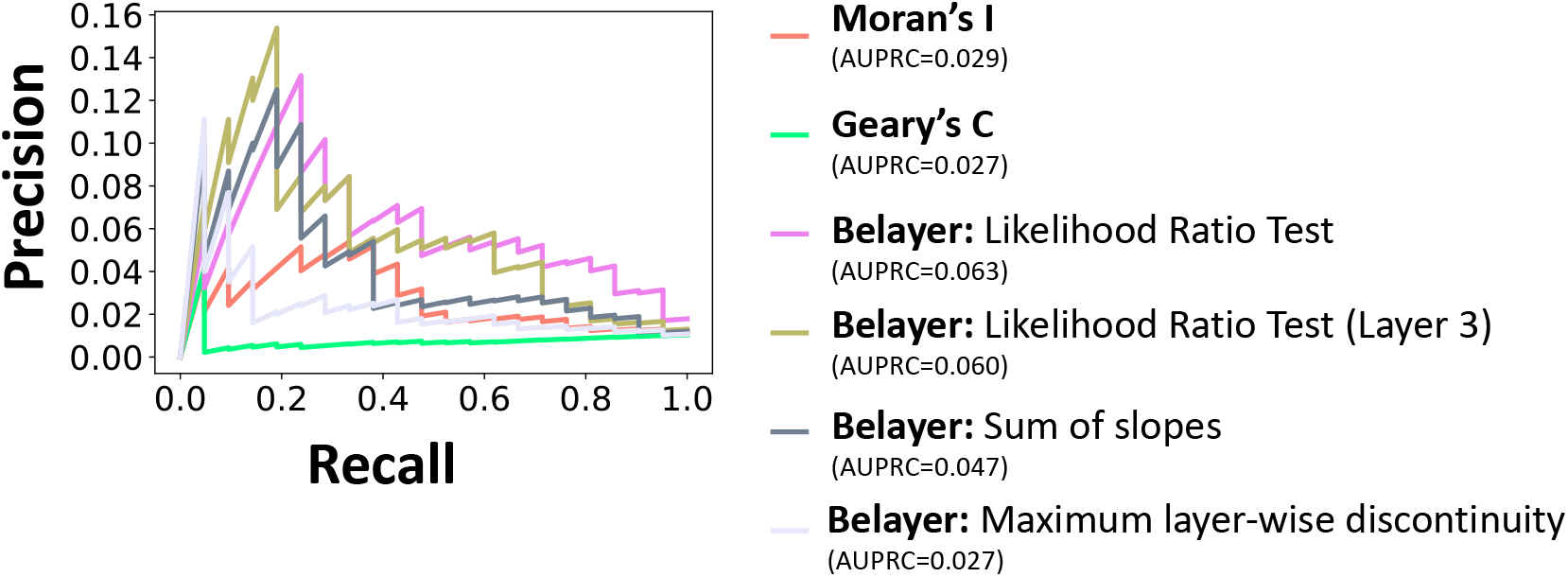
Precision-recall curves for identifying marker genes in DLPFC sample 151508 as in Figure 4A but with different rankings; see Section F for further explanation.

**Figure S11:**
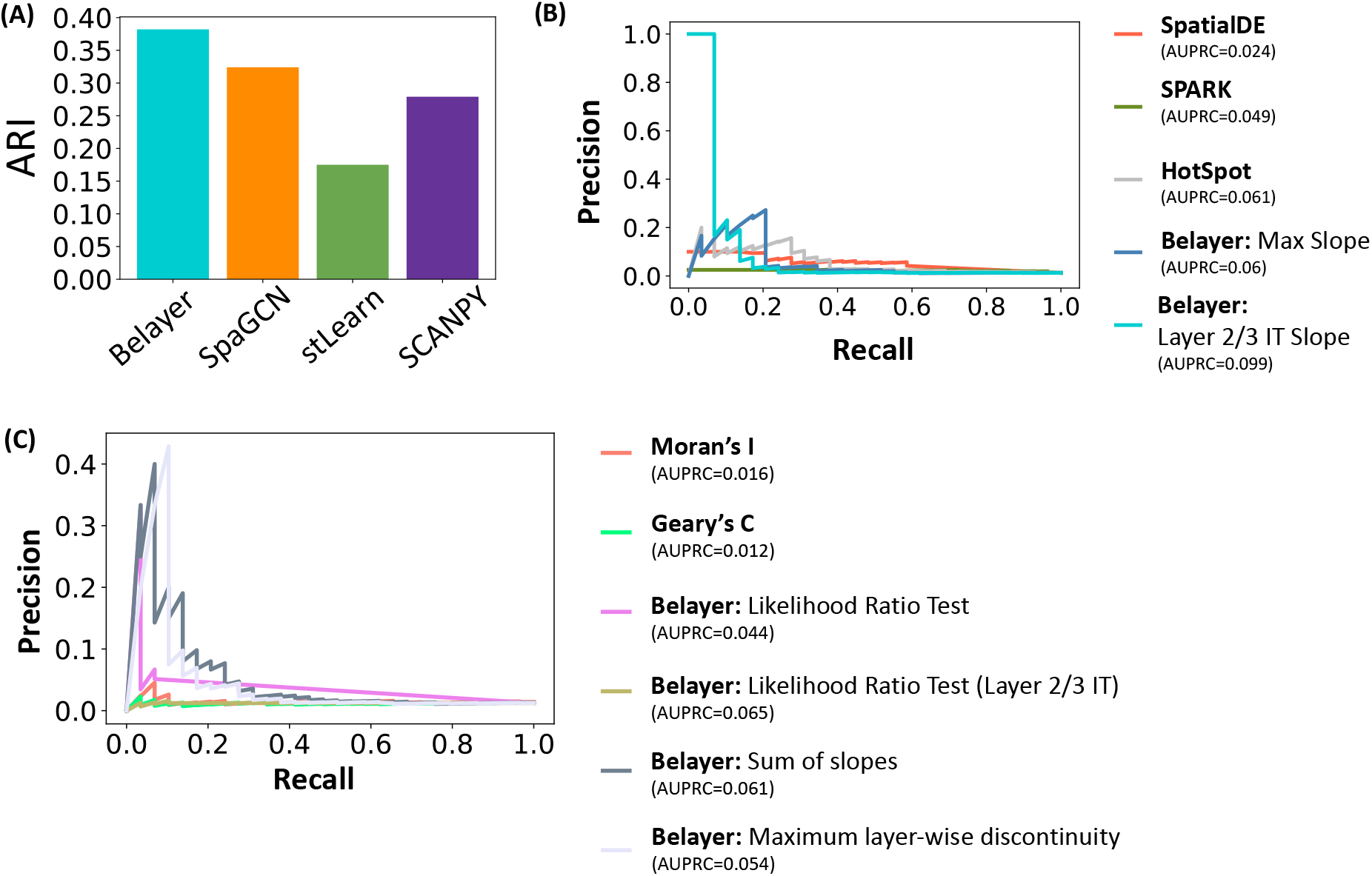
(A) Comparison of Belayer and other methods in identifying cortical layers in Slide-SeqV2 mouse somatosensory dataset. (B)/(C) Precision-recall curves for identifying marker genes in the same dataset using ten different approaches for ranking genes.

**Figure S12:**
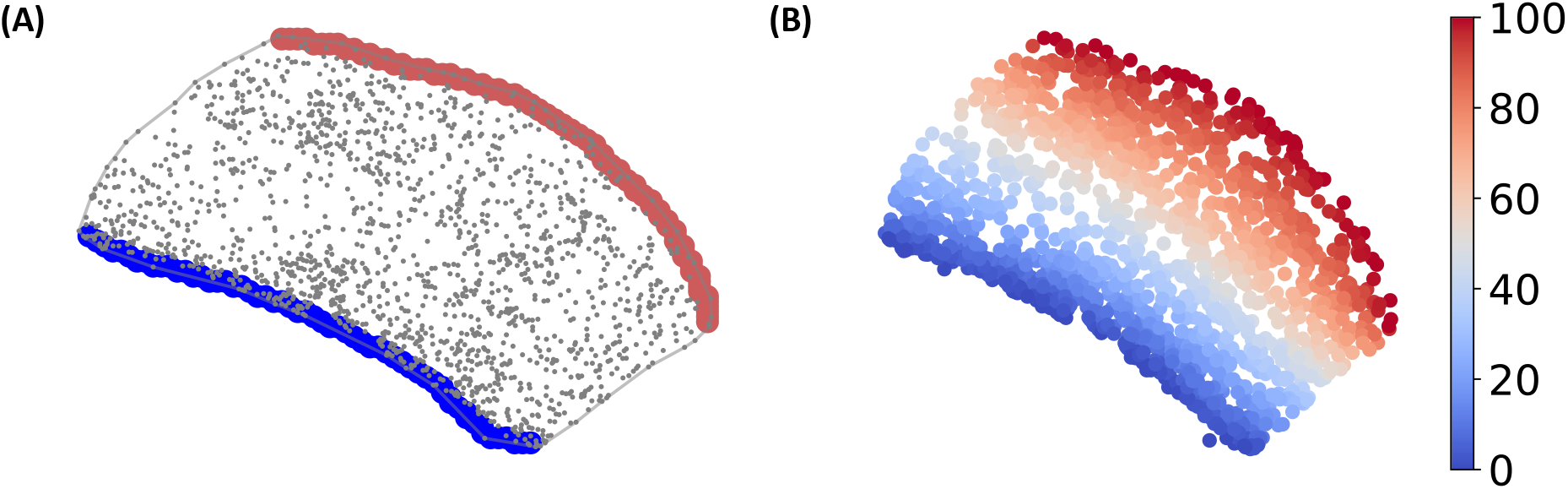
(A) Two parts of the tissue boundary are used to construct a conformal map over the entire tissue in the mouse somatosensory cortex dataset. Each spot is shown in spatial coordinate. Tissue boundary is outlined by a black segmented line; the blue and red points indicate the two involved parts respectively. (B) The real part of the conformal map is equivalent to the solution of heat equation whose values are shown by the colormap.

**Figure S13:**
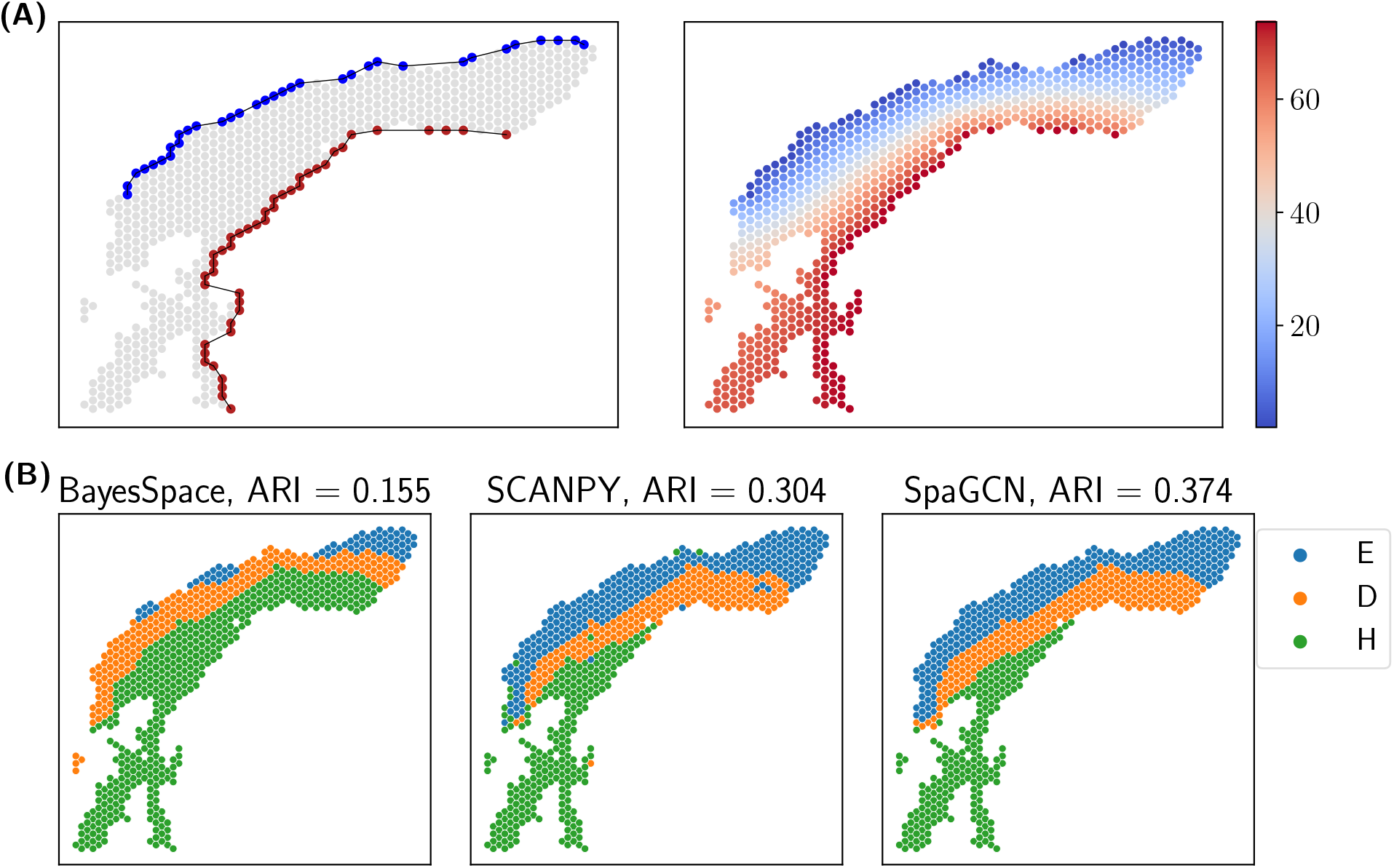
(A) Two parts of the tissue boundary are used to construct a conformal map over the entire tissue (left) in the skin wound dataset. Each spot is shown in spatial coordinate. Tissue boundary is outlined by a black segmented line; the blue and red points indicate the two involved parts respectively. The real part of the conformal map is equivalent to the solution of heat equation whose values are shown by the colormap (right). (B) Clusters identified by three other methods, BayesSpace, SCANPY, and SpaGCN. Clusters are labeled and colored by the maximal overlap with annotated skin layers.

**Figure S14:**
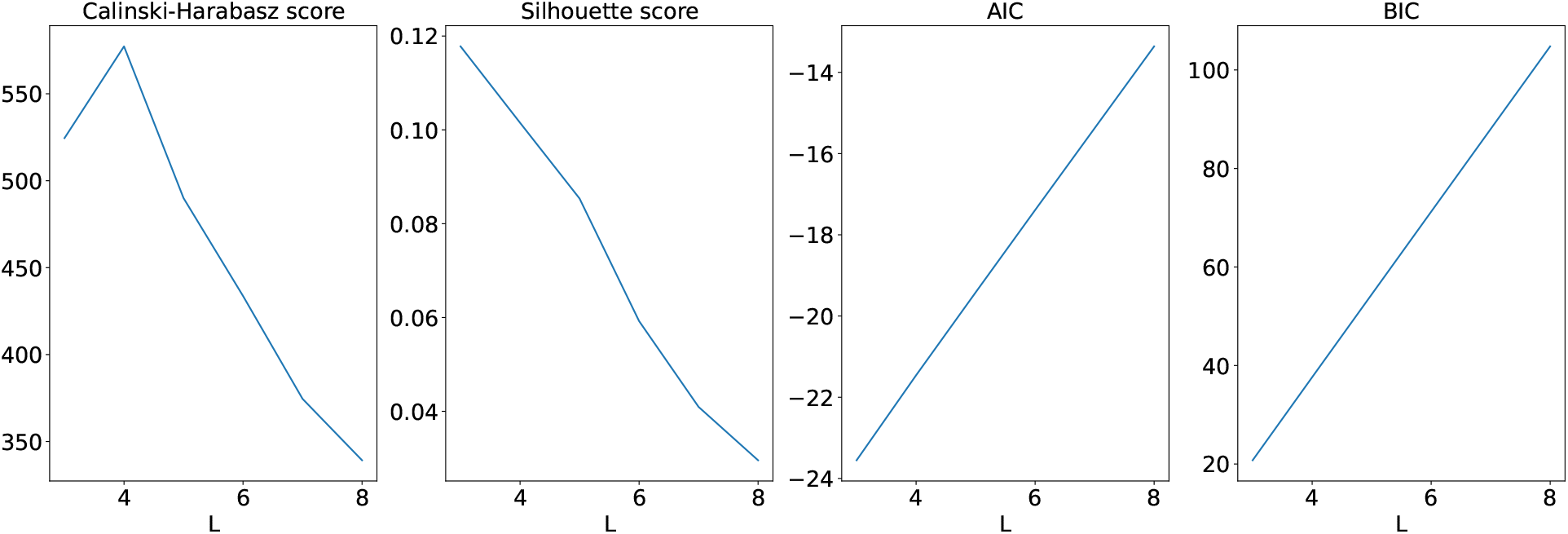
Alternative model selection procedures for choosing the number *L* of layers.

## Notes

### Competing Interest Statement

The authors have declared no competing interest.

### Summary of Updates

Revised mathematical exposition; added new overview Figure; two additional experiments with Slide-SeqV2 mouse somatosensory cortex data and 10x Visium skin wound data

